# Cross-Species Transcriptomic Integration Reveals a MIRO1-Mediated Macrophage–T Cell Axis in Glioma

**DOI:** 10.1101/2025.11.10.686781

**Authors:** Zehui Du, Menghan Li, Brandon H. Bergsneider, Andy P. Tsai, Kwang Bog Cho, Lily H. Kim, John Choi, Gordon Li, Tony Wyss-Coray, Michael Lim, Xinnan Wang

## Abstract

Mitochondrial regulators are increasingly recognized for their influence on immune signaling within the tumor microenvironment (TME). In glioma, where immunosuppression limits therapeutic efficacy, we investigate how targeting the mitochondrial protein MIRO1 alters the TME. We combine single-nucleus RNA sequencing of murine gliomas treated in vivo with a MIRO1-binding compound and bulk RNA sequencing of human glioma resections treated with the same compound ex vivo. Cross-species transcriptomic integration reveals a MIRO1-responsive program in the TME. Among shared targets, we identify *PARP11/Parp11* as a consistently upregulated gene in glioma that is downregulated following MIRO1-binding compound treatment in both human and mouse gliomas. Cell-cell communication analysis shows that a specific cluster of macrophages (MAC1), which exhibits robust *Parp11* and *Pdl1* (encoding PD-L1) expression, sends immunosuppressive signals to CD8+ cytotoxic T cells, and may receive prostaglandin E□ signals from another cluster of macrophages (MAC4). Targeting MIRO1 eliminates this cell circuitry and reduces the tumor cell population. Our study provides a transcriptomic framework for understanding mitochondria-immune crosstalk and nominates MIRO1*-*PARP11 as a potential effector axis of brain immune dysfunction.

**Summary Blurb:** Transcriptomic integration of single-nucleus RNA sequencing of murine gliomas and bulk RNA sequencing of human gliomas reveals an immunosuppressive signaling axis in the glioma TME, mediated by MIRO1.

## Introduction

Glioblastoma (GBM), the highest grade of glioma, is among the most immunosuppressive and treatment-resistant human cancers, in part due to a complex tumor microenvironment (TME) that limits effective anti-tumor immunity. While immune checkpoint inhibitors have transformed outcomes in other malignancies, their efficacy in GBM has been limited, owing to poor T cell infiltration, T cell exhaustion, and strong immunosuppressive cues from tumor-associated myeloid populations, among others (Jackson C et al, 2025, Jackson CM et al, 2024). However, the molecular mechanisms linking myeloid activity to T cell suppression in glioma remain incompletely understood.

MIRO1 (*RHOT1*), an outer mitochondrial membrane (OMM) protein mediating mitochondrial trafficking and quality control (Bharat V et al, 2023, Bharat V et al, 2021, Course MM & Wang X, 2016, Hsieh CH et al, 2019, Hsieh CH et al, 2016, Li L et al, 2021, Nguyen D et al, 2021, Shaltouki A et al, 2018, Vanhauwaert R et al, 2019, Wang X & Schwarz TL, 2009, Wang X et al, 2011), is increasingly recognized for its broader roles in cellular homeostasis (Bharat V & Wang X, 2020, Pekkurnaz G & Wang X, 2022). MIRO1 is inserted into the OMM via a C-terminal transmembrane domain, and its N-terminal cytosolic region contains two EF-hand Ca²□-binding motifs and two GTPase domains. MIRO1’s known functions rely on regulated protein-protein interactions (Glater EE et al, 2006, Hsieh CH et al, 2016, Stowers RS et al, 2002, Wang X & Schwarz TL, 2009, Wang X et al, 2011). We have previously found that MIRO1 protein is upregulated in postmortem brains of sporadic Parkinson’s disease (PD) patients and induced pluripotent stem cell (iPSC)-derived neurons of familial PD patients bearing *SNCA* mutations (Bharat V et al, 2023, Shaltouki A et al, 2018). Stabilization of MIRO1 delays its removal from damaged mitochondria and mitophagy, consequently accumulating oxidative stress and increasing neuronal sensitivity to stressors in PD patient-derived models (Bharat V et al, 2023, Hsieh CH et al, 2019, Hsieh CH et al, 2016, Li L et al, 2021, Nguyen D et al, 2021, Shaltouki A et al, 2018). To therapeutically target MIRO1, we have developed a compound series that bind the C-terminal GTPase domain of human MIRO1 (Hsieh CH et al, 2019, Li L et al, 2021). In PD patient fibroblasts and iPSC-derived neurons, these compounds at the therapeutic dose selectively facilitate proteasomal degradation of MIRO1 upon mitochondrial depolarization, without affecting MIRO1’s overall GTPase activity, other mitochondrial proteins (Bharat V et al, 2023, Hsieh CH et al, 2019, Li L et al, 2021), or basal mitochondrial motility (Hsieh CH et al, 2019). These MIRO1 binders promote mitophagy and alleviate stressor-induced dopaminergic neurodegeneration in PD models (Bharat V et al, 2023, Hsieh CH et al, 2019, Li L et al, 2021).

Beyond PD, targeting MIRO1 has been implicated beneficial in several types of cancer (Boulton DP & Caino MC, 2024, Cangkrama M et al, 2022, Kong F et al, 2025, Nahacka Z et al, 2022, Peng Y et al, 2025, Saha T et al, 2022, Zhou H et al, 2024). Nevertheless, the role of MIRO1 in the glioma TME remains elusive. To explore this question, here we investigate the transcriptional consequences of pharmacologically targeting MIRO1 with a small-molecule MIRO1 binder (MR3) (Bharat V et al, 2023, Hsieh CH et al, 2019, Li L et al, 2021) in the TME of glioma, using two complementary models: single-nucleus RNA sequencing (snRNA-seq) in a mouse model treated in vivo, and bulk RNA-seq of patient’s glioma tissues treated ex vivo. This cross-species approach enables discovery of conserved MIRO1-dependent programs and their effects in the TME.

We identify a transcriptional axis in which MIRO1 activity may drive the expression of *Pdl1* (encoding PD-L1) and *Parp11* in selective macrophages, which could lead to immunosuppression and T cell inhibition in a paracrine manner in the TME. These findings provide new insights into mitochondria-immune crosstalk in glioma and offer a transcriptomic framework for exploring MIRO1 as a modulator of immune suppression.

## Results

### Integration of human and mouse glioma transcriptomic datasets

To investigate how the MIRO1-binding compound (MR3) reshaped glioma transcriptional programs across species, we developed an integrative framework that combined reference tumor gene signatures with drug-treated transcriptomes from both in vivo mouse and ex vivo human models (Fig 1A). As a foundation, we first identified glioma-associated gene expression changes by comparing TCGA (Weinstein JN et al, 2013) glioma samples with non-tumor brain cortex from the GTEx (The genotype-tissue expression (gtex) project, 2013) dataset. This analysis identified a glioma-specific set of transcriptionally dysregulated genes. We used these genes as a baseline to evaluate how MR3 modulated glioma-associated transcriptional programs.

**Figure 1.**
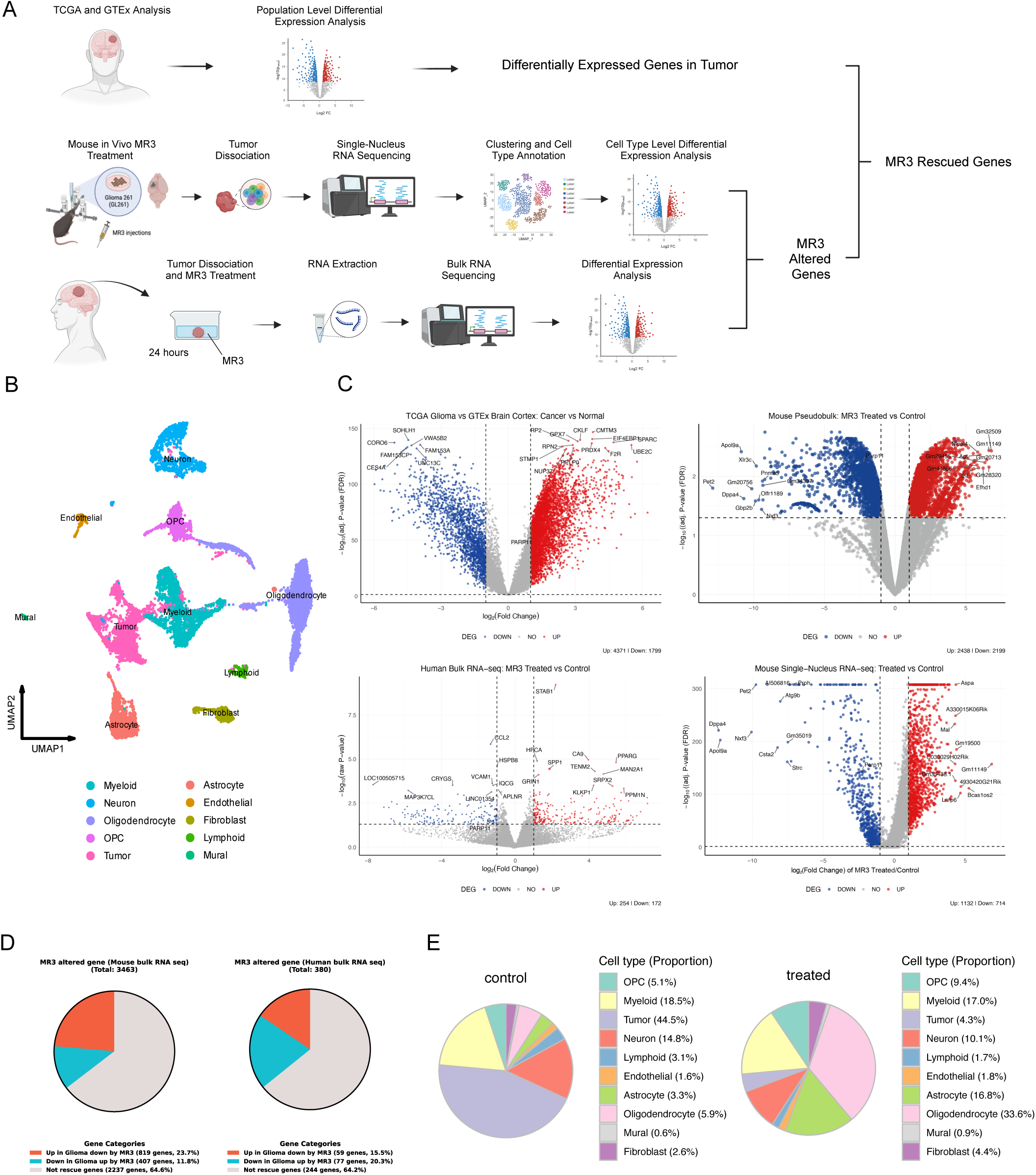
Integration of human and mouse glioma transcriptomic datasets reveals MR3-induced gene expression changes. **(A)** Workflow schematic. Top: Comparison of TCGA glioma and GTEx brain cortex to identify glioma-associated DEGs. Middle: snRNA-seq of GL261 mouse gliomas after MR3 treatment. Bottom: Bulk RNA-seq of resected human gliomas treated ex vivo with MR3 for 24 hours. **(B)** UMAP projection of major cell types in MR3-treated and untreated mouse gliomas, including astrocytes, endothelial cells, fibroblasts, lymphoid cells, mural cells, myeloid cells, neurons, oligodendrocytes, OPCs, and tumor cells. **(C)** Volcano plots of DEGs in TCGA glioma versus GTEx brain cortex (top left), MR3-treated versus untreated (DMSO) human bulk RNA-seq with a paired DESeq2 design including patient as a blocking factor (bottom left), MR3-treated versus untreated (DMSO) mouse glioma pseudo bulk RNA-seq (top right), and MR3-treated versus untreated (DMSO) mouse glioma snRNA-seq (bottom right). Y-axis: −log_10_(raw P-value) for human bulk RNA-seq; −log_10_(adj. P-value (FDR)) for the TCGA/GTEx, mouse pseudo-bulk, and mouse snRNA-seq analyses. Red: upregulated; blue: downregulated; gray: not significant. Significance for the human bulk RNA-seq is defined as raw p < 0.05, whereas significance for the other three comparisons is defined as FDR-adjusted p < 0.05. **(D)** The proportion of MR3-altered genes that are differentially expressed in the opposite direction compared to the TCGA glioma population, in mice (left) and humans (right), calculated within the shared ortholog-mapped gene set. **(E)** Proportions of cell type composition in untreated control (left) and MR3-treated (right) mouse tumors.

To evaluate the impact of MR3 in vivo, we used the syngeneic GL261 mouse glioma model. Mice were injected intracranially with MR3 for the treatment group or the vehicle, DMSO, for the untreated group. Tumor-bearing hemispheres were then collected for snRNA-seq. After quality control, 11,940 NeuN□ nuclei from MR3-treated and untreated mouse samples were integrated using Seurat v5.0 (Butler A et al, 2018, Hao Y et al, 2021, Hao Y et al, 2024, Satija R et al, 2015, Stuart T et al, 2019) and annotated into ten major brain and tumor cell types, including tumor, astrocyte, oligodendrocyte, oligodendrocyte precursor cell (OPC), neuron, myeloid, fibroblast, endothelial, lymphoid, and mural cells (Fig 1B). Data quality was confirmed through standard metrics (Fig S1A-B). Tumor clusters displayed high proliferation and aneuploidy signatures (Fig S1C), and canonical marker expression supported our annotations (Fig S2A-B).

In parallel, we collected freshly resected glioma tissues from three patients with distinct tumor grades (Table S1). Each sample was cultured with the vehicle, DMSO, as a control group, or MR3 as a treatment group for 24 hours before bulk RNA-seq. This approach enabled us to characterize MR3-induced transcriptional responses in human gliomas.

In order to evaluate both tumor-intrinsic and MR3-induced molecular alterations, we next looked at differentially expressed genes (DEGs) across datasets (Fig 1C). TCGA gliomas showed 1,799 downregulated and 4,371 upregulated genes in comparison to GTEx. In the mouse model, pseudo-bulk analysis treating each animal as an independent biological replicate revealed 2,438 upregulated and 2,199 downregulated genes following MR3 treatment, while single-nucleus analysis revealed 1,132 upregulated and 714 downregulated genes. In human samples, paired DESeq2 analysis with patient as a blocking factor identified 254 downregulated and 172 upregulated genes following MR3 treatment. In both species following MR3 treatment, 25 genes were consistently elevated, while 11 genes were downregulated. Despite the relatively modest number of overlapping genes (Fig S2D), the constant directionality across species indicates that MR3 affects a small number of transcriptional targets that are shared by humans and mice. Thus, we were able to successfully combine mouse and human data and identified a conserved group of MR3-responsive genes.

### Impact of MR3 on the TME

We next examined whether MR3 shifted TME gene expression toward a non-tumor state. We focused on genes that were differentially expressed in the MR3-treated mice in the reverse direction relative to the glioma signatures defined by TCGA versus GTEx. In the mouse dataset, 1126 of 3463 MR3-responsive genes (35.4%) showed this reversed pattern. In human tumors, 136 of 380 MR3-responsive genes (35.8%) met the same criterion (Fig 1D). These “rescued” genes may reflect a partial restoration toward non-tumor like expression induced by MR3 treatment.

To evaluate how MR3 reshaped the TME, we analyzed the composition and transcriptomic states of the classified cell types in the mouse snRNA-seq dataset. Sample-level pseudo-bulk PCA showed that MR3-treated mice clustered together and were clearly separated from the control group (Fig S2C), indicating that the transcriptional effect of MR3 exceeded inter-animal variability. MR3 treatment caused clear changes in cell composition (Fig 1E, S2E). Total tumor cell proportion was decreased from 44.5% to 4.3% (Fig 1E), and this reduction was consistently observed in individual MR3-treated animals (Fig S2F). Because macrophages represent one of the most dynamic and treatment-responsive cell populations in the GBM TME (Khan F et al, 2023), we next identified macrophages (described later) in the myeloid population and examined how MR3 treatment changed their gene expression. MR3 treatment showed a clear shift in expression patterns in macrophages (Fig S3A, D). Genes that were increased after treatment were mainly involved in synapse organization, GTPase signaling, cell junction formation, and phosphatidylinositol signaling (Fig S3B, E). These pathways indicate stronger intercellular communication and a more active immune response. Genes that were decreased were linked to viral response, type I interferon signaling, collagen structure, and MHC class I peptide loading (Fig S3C, F). Together, the data show that MR3 pushed macrophages toward a more active, immune-supporting state.

We then performed Gene Ontology (GO) enrichment analysis on DEGs in bulk RNA-seq data from MR3-treated and untreated patient glioma samples (Fig 2A). Compared with the untreated control group, the treated samples showed upregulation of immune activation-related pathways, including chemotaxis, leukocyte activation, positive regulation of cell adhesion, αβ T cell activation, and immune response-activating cell surface receptor signaling. No significantly enriched downregulated biological processes were detected. These findings suggest that MR3 may enhance anti-tumor immune responses.

**Figure 2.**
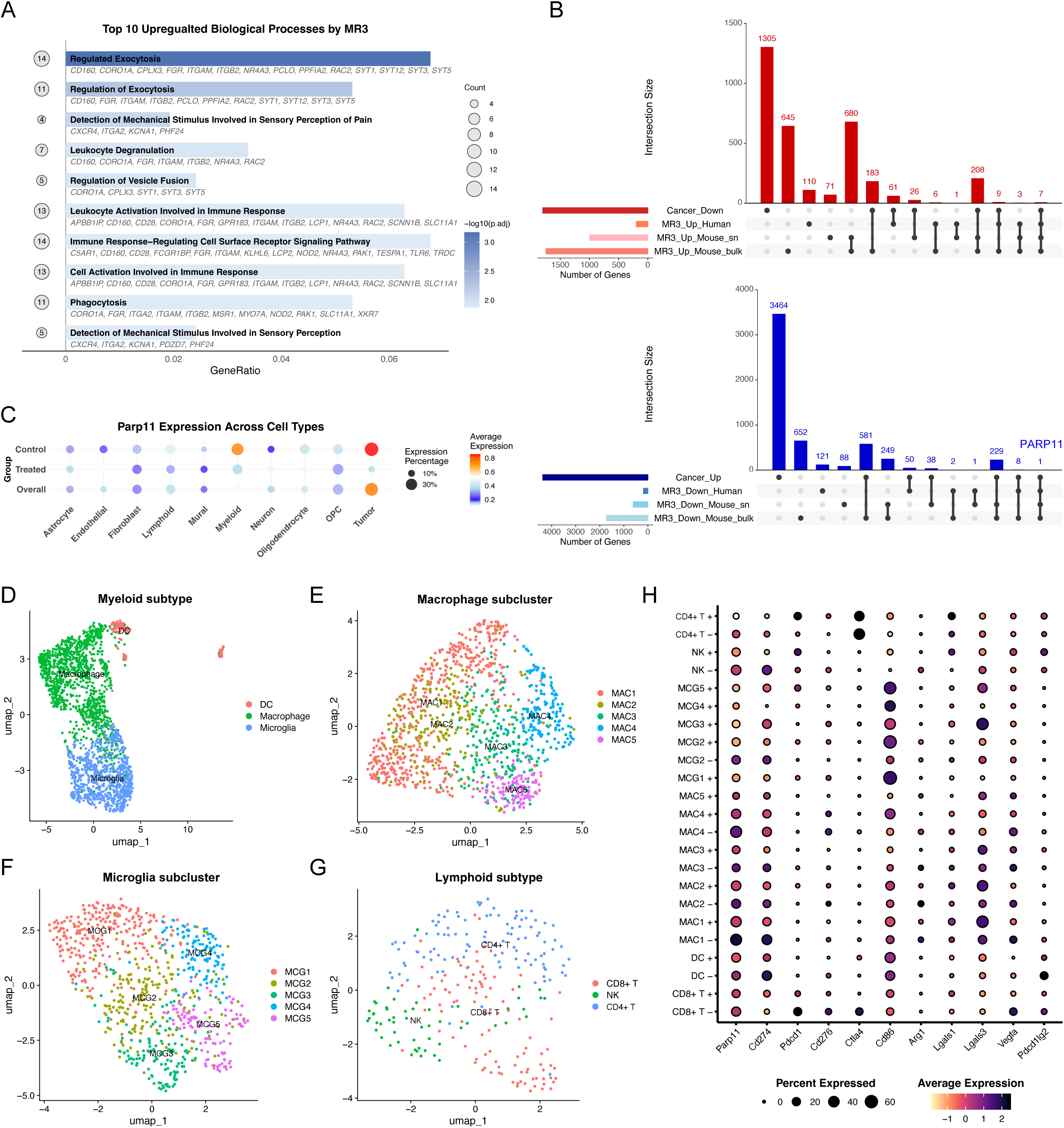
Cross-species transcriptomic analysis identifies *PARP11/Parp11* as a conserved MIRO1-responsive gene. **(A)** GO enrichment analysis of DEGs from human bulk RNA-seq after MR3 treatment, showing the top 10 significantly upregulated biological processes. **(B)** UpSet plots comparing glioma-dysregulated genes from TCGA (cancer vs. GTEx cortex) with MR3-regulated genes across mouse snRNA-seq, mouse pseudo-bulk RNA-seq, and human bulk RNA-seq. Among cancer-upregulated genes, *PARP11/Parp11* emerged as the only gene consistently downregulated by MR3 in all three analyses. **(C)** Dot plots depicting the expression of *Parp11* across all cell types in untreated control and treated conditions. Dot size indicates the proportion of cells expressing each gene, and color intensity reflects the average expression level. **(D-G)** UMAP projections showing myeloid subclusters: macrophage, microglia, and DCs (D); macrophage subclusters: MAC1-MAC5 (E); microglia subclusters: MCG1-MCG5 (F); lymphoid subclusters CD8□ T, NK, and regulatory-like CD4□ T cells (G). **(H)** Dot plot showing the expression of *Parp11* and other immune suppression genes in immune cell subclusters. +: MR3; -: no MR3.

### *PARP11/Parp11* is a conserved MIRO1-responsive gene in the glioma TME

To identify key downstream targets of the MIRO1 signaling, we compared glioma-dysregulated genes from TCGA (cancer vs. GTEx normal cortex) with MR3-induced gene expression changes across three analyses: mouse snRNA-seq, mouse pseudo-bulk from snRNA-seq, and human bulk RNA-seq. We asked whether MR3 could reverse glioma-associated transcriptional patterns and rescue a subset of cancer-dysregulated genes.

In the mouse model, we found 819 downregulated genes in the pseudo-bulk analysis and 268 in the single-nucleus data by MR3 overlapped with cancer-upregulated genes. Their intersection yielded 229 candidates. In the human dataset, 59 cancer-upregulated genes were downregulated by MR3. Only one gene, *PARP11/Parp11*, was consistently downregulated by MR3 across all three analyses and simultaneously upregulated in glioma cancer dataset (Fig 2B). Conversely, seven genes (*ADAM11*, *KCNA1*, *PAK1*, *PPFIA2*, *SLC24A4*, *SNX32*, and *SYNPR*) were consistently upregulated by MR3 and downregulated in cancer. Given the reported role of *PARP11/Parp11* in regulating tumor immunosuppression and its association with response to immune checkpoint blockade (Basavaraja R et al, 2024, Bisht P et al, 2022, Wang S et al, 2024, Zhang H et al, 2022), we prioritized *PARP11/Parp11* for further investigation.

We next asked where *Parp11* was expressed and how its levels were changed after treatment in the mouse snRNA-seq dataset. *Parp11* showed high expression in both tumor and myeloid cells, and MR3 treatment led to a clear reduction in both average expression and the proportion of expressing cells in these compartments (Fig 2C).

Interestingly, past research has shown that PARP11 promotes the immunosuppressive function of tumor-infiltrating regulatory T cells (Tregs): its expression correlates with poor response to immune checkpoint blockade, and pharmacological inhibition of PARP11 can reactivate intra-tumoral T cells and enhance the efficacy of checkpoint inhibitor and CAR T cell therapies (Basavaraja R et al, 2024, Wang S et al, 2024, Zhang H et al, 2022). Across multiple cancer types, *PARP11* expression correlates with patient survival during anti-PD-1 treatment (Wang S et al, 2024). In GBM, PARP activities have been linked to chemoresistance, and PARP inhibitors are being explored as adjuncts to temozolomide therapy (Bisht P et al, 2022). This evidence supports the immune-related responses by MR3 treatment in our datasets. Together, we identify *PARP11/Parp11* as a conserved MIRO1-dependent gene that is consistently suppressed by MR3 across species and various datasets.

### MAC1 is enriched with *Parp11* and *Pdl1* expression

To investigate the mechanistic link between *Parp11* and MR3, we further subclustered immune-related cells and tumor cells in the mouse snRNA-seq dataset (Fig S3G). After batch effect correction, the myeloid compartment fell into three major lineages defined by canonical markers: *Ms4a7* for macrophages, *P2ry12* for microglia, and *H2-Ab1* for dendritic cells (DCs) (Fig 2D, S4A). Within the macrophage lineage, we identified five subpopulations (MAC1–MAC5) based on DEGs (Fig 2E, S4B, G). Microglia were subdivided into five clusters (MCG1–MCG5) (Fig 2F, S4B, G). Lymphoid subset was categorized as CD8□ T cells (*Cd8a*), regulatory-like CD4□ T cells (*Il2ra*), and natural killer (NK)-like cells (*Klrk1*) (Fig 2G, S4C-D, G). Tumor cells were also divided into five transcriptional states (T1–T5) based on distinct expression patterns (Fig S4E-F, H).

To resolve the immunosuppressive components more precisely, we examined *Parp11* and other immune suppression-related genes across the myeloid and lymphoid subclusters. We found that *Parp11* was most enriched in MAC4 and MAC1 (Fig 2H). MR3 treatment reduced both *Parp11* expression level and the fraction of expressing cells in these subsets (Fig 2H).

When we examined other genes linked to immune suppression, we found that MAC1 also showed high *Cd274* (*Pdl1*, encoding PD-L1) levels, a well-known marker of immune escape. After MR3 treatment, *Cd274* levels were decreased, similar to *Parp11* (Fig 2H). This pattern suggests that MR3 may weaken immunosuppressive signals by lowering both PARP11 and PD-L1 expression in MAC1 cells.

### Cell-cell communication analysis suggests macrophage-derived signaling drives *Parp11* upregulation and T cell inhibition

Having identified MAC1 as a macrophage population enriched for *Parp11* and *Pdl1*, we next performed cell-cell communication analysis to explore its downstream targets. Among all cell-cell signaling and communication pathways, the overall number and strength of interactions increased after MR3 treatment (Fig S5A-E).

Next, we analyzed MAC1 as the signal sender and examined all outgoing interactions originating from this cluster. The overall interaction from MAC1 to lymphoid cells was increased after MR3 treatment (Fig 3A, B). Among these changes, we observed the elimination of immunosuppressive pathways targeting CD8+ T cells, including *Cd274-Pdcd1* (PD-L1/PD-1) (Stuart T et al, 2019, Topalian SL et al, 2015) and *Cd86-Ctla4* (Ha D et al, 2019, Qureshi OS et al, 2011). In contrast, immune-activating signals like *Cd86-Cd28* (Fuse S et al, 2006) were enhanced. Consistent with this shift in predicted intercellular communication, we observed upregulated expression of multiple genes associated with CD8□ T cell activation, cytotoxic function, and interferon response following MR3 treatment (Fig S6A). These findings suggest that MR3 may suppress immunoinhibitory communication while activating pro-immune signaling, potentially restoring CD8□ T cell activity.

**Figure 3.**
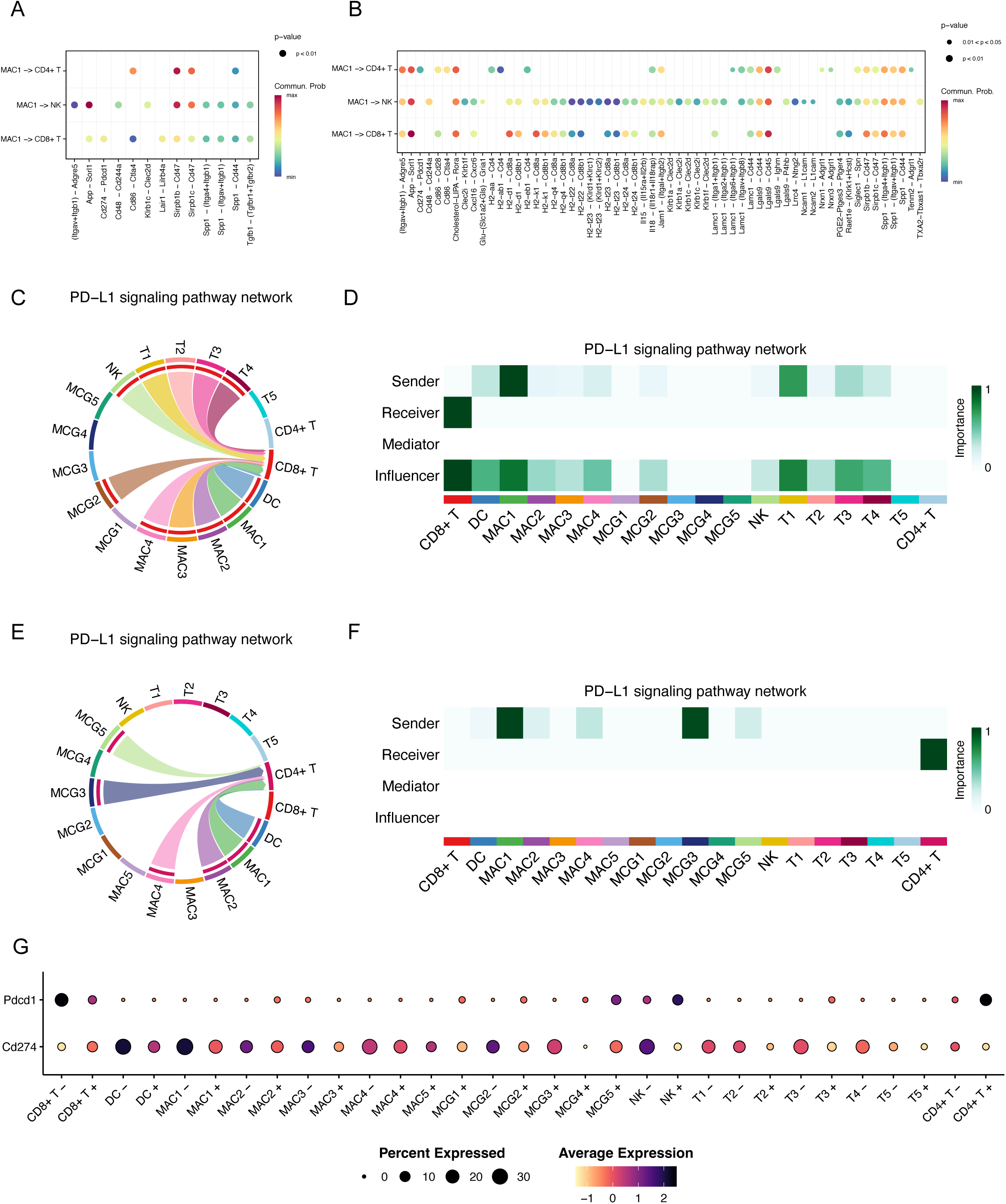
Cell-cell communication analysis reveals MR3-dependent changes in intercellular communication. (A, **B)** Dot plots depicting CellChat analysis of cell-cell communication in the MR3□ (A) and MR3□ (B) conditions, with MAC1 as the signaling source and interactions directed toward other clusters. Dot size represents significance, and color indicates communication probability. Subclusters with fewer than 20 cells per condition are filtered out, which may result in some subclusters being absent in certain conditions. These plots represent predicted signaling probabilities inferred from transcriptomic data, and do not measure functional interactions. **(C, E)** Chord plot illustrating CellChat analysis of PD-L1/PD-1 signaling interactions among subclusters in (C) MR3□ and (E) MR3^+^ condition. Subclusters with fewer than 20 cells per condition are not shown due to filtering. The chord plots indicate predicted ligand-receptor communication probabilities rather than direct protein-level interactions. **(D, F)** Heatmap demonstrating network centrality measures for the PD-L1/PD-1 signaling network. The metrics indicate the relative importance of each subcluster in sending, influencing, and receiving PD-L1/PD-1 signals. (D) MR3□ and (F) MR3^+^. Subclusters with fewer than 20 cells per condition are omitted, and the centrality measures reflect predicted signaling roles rather than functional activity. **(G)** Dot plot showing the expression of *Cd274 (Pdl1)* and *Pdcd1 (Pd1)* across immune cells and tumor subclusters in MR3□ and MR3□ conditions. Subclusters with fewer than 20 cells are not shown. These expression levels support predicted signaling but do not demonstrate direct inhibitory effects.

We next specifically examined PD-L1/PD-1 signaling in MR3-treated and untreated groups across all cell subclusters. In untreated controls, MAC1 was consistently identified as the main source of *Cd274* (*Pdl1*) signals directed toward CD8□ T cells via the *Pdcd1* (*Pd1*, encoding PD-1) receptor (Fig 3C, D). After MR3 treatment, this interaction was substantially reduced (Fig 3B, E, F): MAC1 no longer sent PD-L1/PD-1 signals to CD8□ T cells. In contrast, regulatory-like CD4□ T became recipients of the PD-L1/PD-1 signaling axis, receiving signals mainly from MAC1 and MCG3 (Fig 3E, F). This may be due to compensatory upregulation of *Pd1* in regulatory-like CD4□ T in response to decreased PD-L1 in the TME (Fig 3G), consistent with previous reports showing that disruption of PD-1/PD-L1 signaling can induce compensatory expression of inhibitory receptors on T cells (Huang RY et al, 2017, Saleh R et al, 2019).

Tumor clusters did not exhibit significant outgoing PD-L1/PD-1 signaling to CD8□ T cells (Fig 3D), despite high *Parp11* expression (Fig 2C). This suggests that tumor cells play a minimal role in mediating PD-L1/PD-1-dependent suppression in this model, and MAC1 is the primary source of T cell inhibition via this axis.

We then asked what upstream mechanisms sustained *Parp11* expression in MAC1. Using the current CellChatDB.mouse database, we detected several signaling pathways upstream of MAC1 (in which MAC1 was the receiver of signals) that were altered after treatment; however, none of these interactions were shown by previous work or our data to be associated with the induction of PARP11, MIRO1, or MR3 (Fig S5F, G).

Next, we probed individual gene expression patterns for possible mechanisms. Previous studies showed that PARP11 expression can be induced by interferon signaling (Guo T et al, 2019, Lüscher B et al) or the cAMP-PKA-CREB pathway (Basavaraja R et al, 2024). We therefore examined expression of their respective relevant genes: type I interferon (IFN-I)-stimulated genes (ISGs) and prostaglandin E□ (PGE□) biosynthesis genes, across all myeloid clusters (Fig 4A).

**Figure 4.**
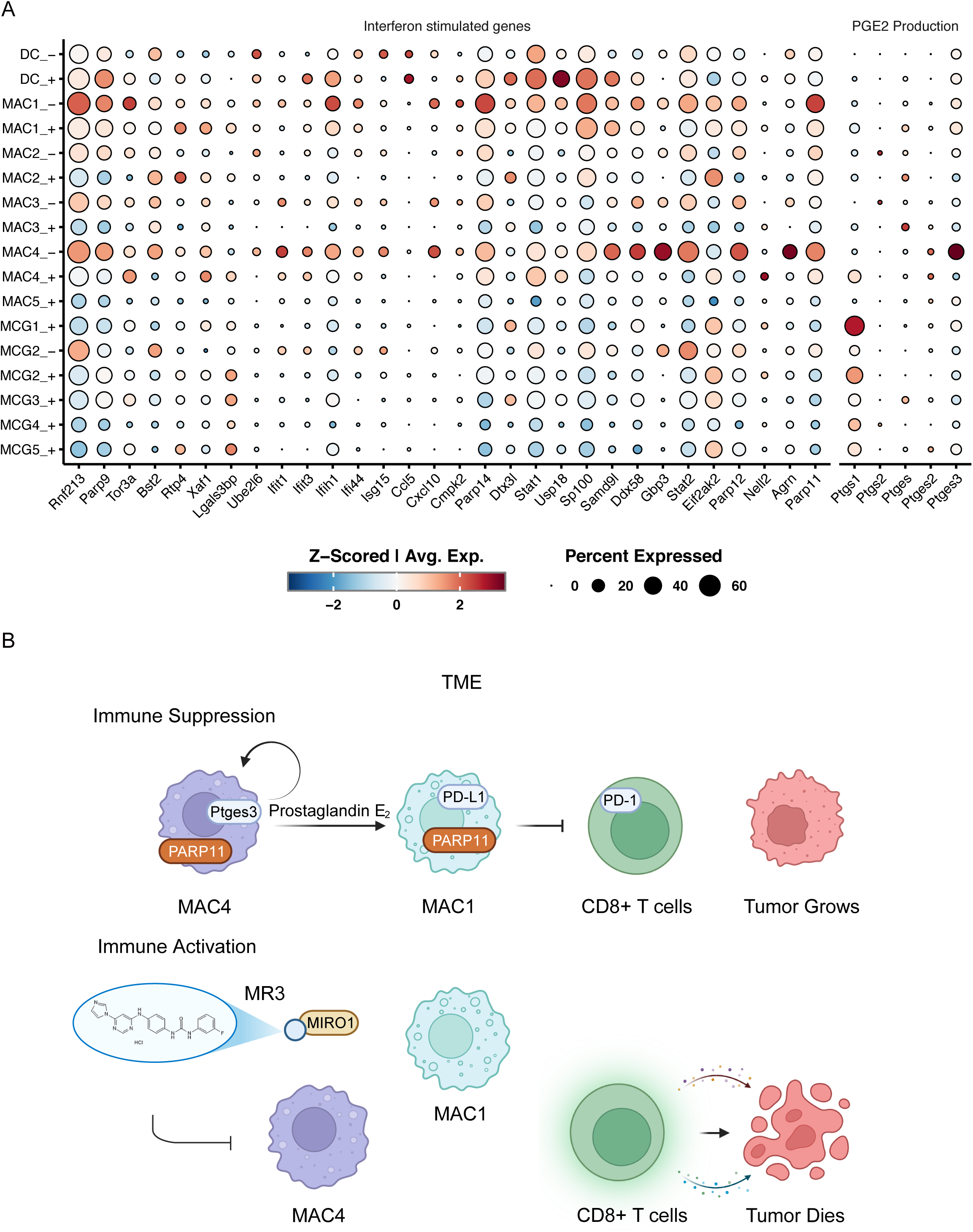
MR3 treatment alters myeloid gene expression and reshapes the TME. **(A)** Dot plot showing the expression of ISGs and PGE_2_ production-related genes across myeloid subclusters in MR3□ and MR3□ conditions. The size of each dot represents the proportion of cells expressing the gene, and the color intensity indicates the average expression level. Subclusters with fewer than 20 cells were excluded. **(B)** Schematic model illustrating how MR3 treatment reshapes the TME.

Mitochondria stress-triggered ISGs, such as *Ifit1*, *Ifit3*, and *Isg15*, were highly expressed in both MAC4 and MAC1 untreated control groups, suggesting active mitochondrial stress or an IFN-I response (Baldanta S et al, 2017, Boccuni L et al, 2022). MR3 treatment reduced the expression of these genes (Fig 4A). However, we did not detect IFN-I transcripts directly, so the exact producing subcluster remained unknown. Even though the source could not be clearly defined, the consistent drop in ISG expression after MR3 treatment suggests a weakened IFN-I-driven program, and IFN-I may still play a role in maintaining *Parp11* expression in MAC1.

Among genes related to prostaglandin production, *Ptges3*, encoding PTGES3, the terminal synthase responsible for PGE□ formation, was found to be highly expressed only in the MAC4 untreated group and was sharply reduced after MR3 treatment (Fig 4A). Previous studies showed that PGE□ can activate the PKA-CREB pathway, which induces PARP11 expression (Basavaraja R et al, 2024). Intriguingly, PGE□ may contribute to the induction of myeloid-derived suppressor cells and immunosuppression in glioma (Dean PT & Hooks SB, 2022, Mi Y et al, 2020). Together, these bioinformatic findings suggest that MAC4 may produce PGE□, which could act on nearby MAC1 cells in a paracrine manner to increase *Parp11* expression, although this model needs to be functionally validated.

### MIRO1-related pathways show modest changes by MR3

To further understand the mechanistic effects of MR3, we screened several other MIRO1-relevant signaling modules, including NF-κB, mitochondrial motility, dynamics, mitophagy, the cGAS–STING axis, and mitochondrial DNA (mtDNA) release-associated pathways. We found no substantial differences in the expression of NF-κB-related genes in either MAC4 or MAC1 upon treatment (Fig S6B). Genes associated with mitochondrial dynamics, motility, or mitophagy were increased slightly after MR3, most notably in microglia, DCs, or MAC1 (Fig S6C), suggesting a modest enhancement of mitochondrial quality control, consistent with a pro-mitophagy role for MR3 identified by us previously (Bharat V et al, 2021, Hsieh CH et al, 2019, Li L et al, 2021). In MAC1, genes linked to mtDNA release and the cGAS-STING-IRF3 pathway were increased after MR3 treatment (Fig S6D), indicating the involvement of innate immune pathways, which requires future functional investigation.

### A possible paracrine model in glioma TME

Based on these findings, we propose a possible mechanism of MIRO1-mediated immunosuppression in glioma (Fig 4B). In untreated glioma, MAC4 highly expresses *Ptges3*, which produces PGE□ and releases it into the TME. PGE□ then acts on MAC1 in a paracrine manner and activates the cAMP-PKA-CREB pathway, which induces *Parp11*, promoting an immunosuppressive phenotype. Through PD-L1/PD-1 signaling, MAC1 suppresses CD8□ T cells, inhibiting their ability to kill tumor cells. MIRO1 might contribute to this immunosuppressive circuitry by aberrant protein-protein interactions in certain cell clusters and MR3 disables those gain-of-function interactions. As a consequence, *Ptges3* in MAC4 is reduced leading to the decrease of PGE□ in MAC4 and reduction of PARP11 and PD-L1 in MAC1. Ultimately, CD8□ T cells are no longer suppressed by the PD-L1 signal and can restore their ability to kill tumor cells. This model is based on our bioinformatic analysis and warrants further validation in functional studies.

### MiroScape: an interactive portal for cross-species glioma transcriptomics

To enhance accessibility and facilitate broader hypothesis generation, we created MiroScape, a publicly available online interactive web platform (https://miroscape.github.io/MiroScape/). It integrates multiple levels of MIRO1-related datasets from our lab’s research, including transcriptomic, mitochondrial surface proteomic, and proteomic data. The data from this study are available under the MiroScripts module, which hosts the cross-species transcriptomic analyses generated in this work (Fig 5A).

**Figure 5.**
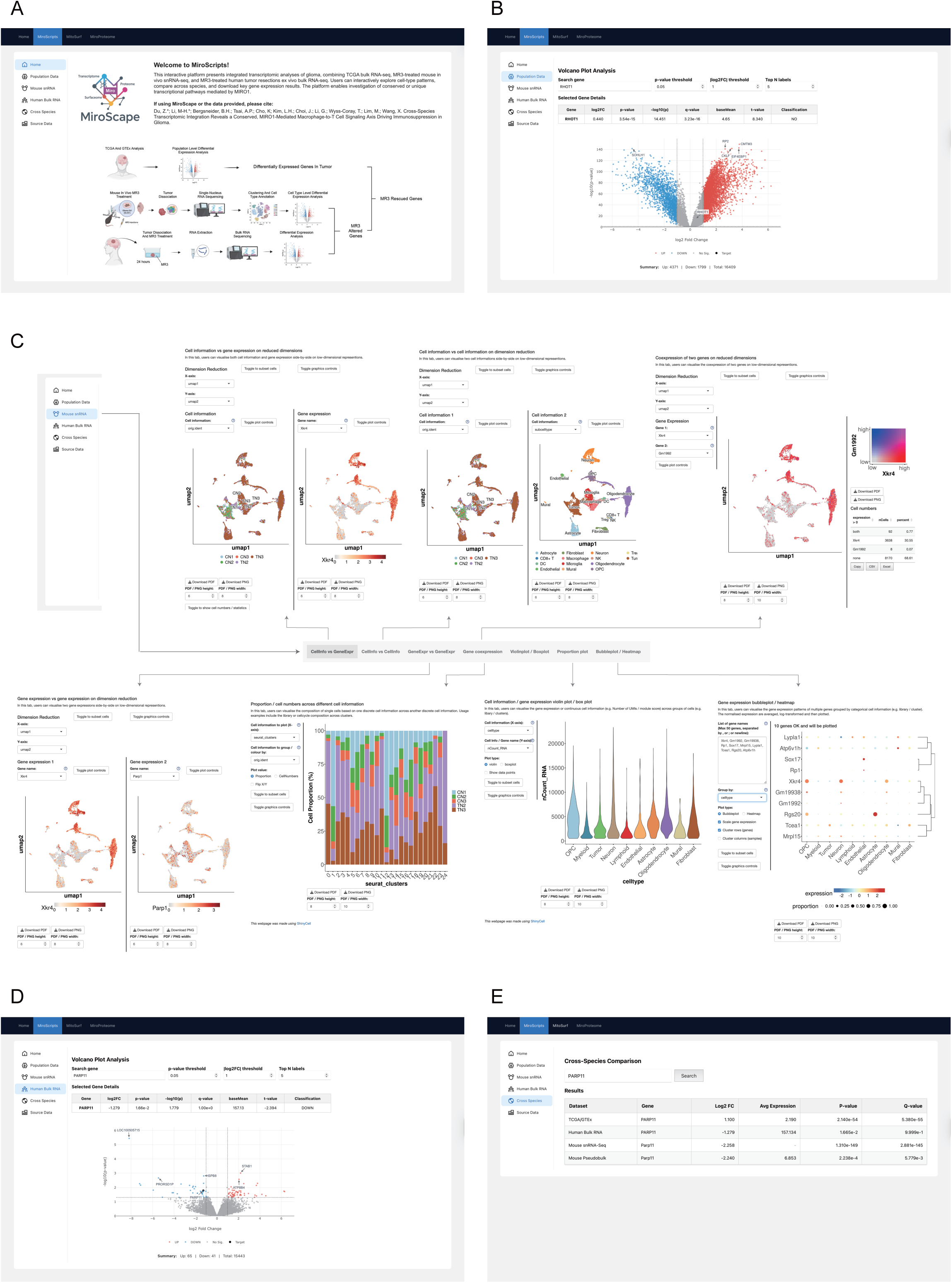
MiroScape website. **(A-E)** Screenshots of MiroScape. (A) MiroScripts home page. (B) Population Data module. (C) Mouse snRNA module featuring CellInfo vs GeneExpr, CellInfo vs CellInfo, GeneExpr vs GeneExpr, Gene coexpression, Violinplot / Boxplot, Proportion plot, and Bubbleplot / Heatmap. (D) Human Bulk RNA module. (E) Cross Species module.

In the Population Data tab (Fig 5B), we provide differential expression results comparing glioma samples from TCGA to non-tumor cortex samples from GTEx. Users can search any gene of interest for its log□ fold change, average expression, t-statistic, p-value, and q-value to identify transcriptional dysregulation in glioma patients. In the Mouse snRNA module (Fig 5C), we present snRNA-seq data from MR3-treated and untreated control mouse glioma samples. We use the ShinyCell2 engine to support interactive visualization. Users can explore annotated cell types, look at gene expression across clusters, and access detailed cell-level metadata. This enables investigation of how different brain and tumor cell populations respond to MR3 in vivo. In the Human Bulk RNA module (Fig 5D), we offer bulk RNA-seq results comparing MR3-treated and untreated glioma samples from patients. As in the Population Data module, users can search for genes and retrieve log□ fold change, average expression, and related statistics. This provides insight into the effects of MR3 in human tissue ex vivo. To connect results across species and sequencing platforms, we designed a Cross Species module (Fig 5E). This feature allows users to view MR3-responsive genes from human and mouse datasets side by side and to recognize shared transcriptional patterns and conserved targets. Users can also explore the full list of differentially expressed genes and download the original data in the Source Data module for deeper analysis. Our web platform is built to help researchers navigate data flexibly and interactively, providing a practical resource for future studies on tumor immunology and therapy development.

## Discussion

Our cross-species transcriptomic analysis uncovers a MIRO1-dependent immune axis in glioma in which certain macrophages may release effectors to the TME, likely inducing increased expression of PARP11 in selective macrophage cell clusters and leading to T cell inhibition and immunosuppression in a paracrine manner. Notably, a MIRO1-binding compound specifically disables this axis, reversing immunosuppressive signatures in the TME, indicating the involvement of mitochondria in driving macrophage immunosuppressive phenotypes.

PARP11, a less characterized member of the PARP family, is emerging as a regulator of post-translational ADP-ribosylation, with possible roles in antiviral signaling and immune modulation (Basavaraja R et al, 2024, Wang S et al, 2024, Zhang H et al, 2022). Its conservation across both species and selective expression within a macrophage subset (MAC1), which may respond to signals from another macrophage cluster (MAC4), highlights PARP11 as a candidate mediator of T cell suppression downstream of mitochondrial signaling cues. The observed variability in treatment response is consistent with the well-established molecular and phenotypic heterogeneity of human glioma. Importantly, despite this diversity, *PARP11/Parp11* is consistently altered across all patient’s samples and the mouse model. This cross-species reproducibility supports its biological and translational relevance.

In parallel, we have found that the originating MAC4 cluster displays robust MIRO1-dependent *Ptges3* expression. *Ptges3* could contribute to PGE□ production and release into the TME, inducing *Parp11* expression in both MAC4 and MAC1 clusters (Fig 3, 4A-B) (Basavaraja R et al, 2024, Wang S et al, 2024, Zhang H et al, 2022). The results raise the possibility that MIRO1 contributes to processes other than its known function in mitochondrial transport and mitophagy. MIRO1 may also act as a regulator of immune signaling within the TME. Previous studies have shown that MIRO1 is required for mtDNA leak into the cytosol in several cell types (Konig T et al, 2021, Nguyen M et al, 2025). Such leakage may activate innate immune pathways through the cGAS-STING signaling (Newman LE et al, 2024, Platanias LC, 2005, Victorelli S et al, 2023, West AP et al, 2015, Zecchini V et al, 2023), linking mitochondrial stress to immune activity in tumors. Thus, it is possible that MIRO1 mediates *Ptges3* activation in this specific cluster of macrophages via mtDNA-stimulated signaling. The resulting paracrine model, in which MIRO1-regulated myeloid states influence T cell gene expression, offers a new perspective on how mitochondrial regulators shape immune dynamics in glioma.

Although our study is transcriptomic in nature and lacks genetic *MIRO1* perturbations, the consistent cross-species findings, context-specific gene programs, and prior validation of the specificity of MR3 (Hsieh CH et al, 2019, Li L et al, 2021), support the biological relevance of this axis. The macrophage cell clusters enriched for PARP11 warrant further investigation to determine their functional phenotypes and potential sensitivity to combined MIRO1 and PARP11 inhibition. A new direction of research is to investigate cellular and immune functions of MIRO1 in macrophages to better understand mitochondria-immune crosstalk in the TME.

Altogether, our work provides a transcriptomic resource and systems-level model of mitochondrial regulation of immune suppression in glioma. MIRO1*-*PARP11 may act as a potential effector axis linking myeloid cell reprogramming to T cell dysfunction in the TME.

## Materials and Methods

### TCGA and GTEx Comparison Analysis

We analyzed population-level transcriptomic changes between glioma and non-tumor cortex using GEPIA3 (Kang YJ et al, 2025) (https://gepia3.bioinfoliu.com/), which integrates TCGA and GTEx RNA-seq datasets. The analysis compared RNA-seq profiles from 166 glioma samples in TCGA with 110 non-tumor brain samples, including 105 cortex samples from GTEx and 5 peritumoral samples from TCGA. We restricted the analysis to protein-coding genes and applied the limma-based statistical framework implemented in GEPIA3. Genes with FDR-adjusted p-values below 0.05 and an absolute log□ fold change greater than 1 were considered significantly differentially expressed.

### Intracranial Glioma Implantation and Monitoring

Although GL261 model demonstrates higher immunogenicity compared to human GBM, this feature enables evaluation of immune-modulating therapies and mechanisms in an immune-competent setting. This model preserves critical aspects of glioma biology, including immunosuppressive TME, invasive behavior, and intracranial growth (Mathios D et al, 2016). Female C57BL/6J mice (6-8 weeks) were obtained from the Jackson Laboratory and housed at the Stanford University Animal Facility under an IACUC protocol. Mice were anesthetized with ketamine (100 mg/kg) and xylazine (10 mg/kg) via i.p. injection. Topical eye gel was applied for lubrication. To establish a syngeneic glioma model, a midline incision was made to expose the skull. A burr hole was drilled over the left striatum (2 mm lateral and 2 mm posterior to the sagittal and coronal sutures), and 1.3 × 10□ GL261-Luc cells (2 µl in PBS) were stereotactically implanted at a depth of 3 mm from the cortical surface. Tumor growth was monitored on day 7 using an IVIS platform (Lago, Spectral Instrument Imaging) following i.p. injection of 200 µL D-luciferin (150 mg/kg for a 20 g mouse; LUCK, Gold Biotechnology). Mice were assigned to 0 µM or 10 µM treatment groups based on luminescence intensity to ensure comparable tumor burden across groups.

### MR3 Treatment and Tissue Collection

Three mice received intracranial administration of MR3 in 5 µL DMSO (10 µM) at the tumor implantation site on days 11 and 18 post-implantation, and three mice received 5 µL of DMSO only at the same time. All intracranial injections followed the same surgical procedure used for tumor implantation. All mice were euthanized on day 22. Whole brains were snap-frozen in liquid nitrogen and stored at –80°C. Only the bottom region of the left hemisphere, where the tumor was implanted, was used for snRNA-seq.

### Nuclei Isolation and Fluorescence-Activated Cell Sorting (FACS)

Nuclei were isolated from the fresh-frozen brain samples of mice using Dounce homogenizers (D8938, Sigma-Aldrich) and the Nuclei EZ Prep Kit (Nuc101-1KT, Sigma-Aldrich) according to previously published methods (Tsai AP et al, 2023). The nuclei were then centrifuged at 300 g for 10 minutes at 4°C and resuspended in 50 μL of FACS buffer (1% BSA, 1×PBS; sterile filtered) containing 2 U/ml of Recombinant RNase Inhibitor (2313B, Takara) and anti-CD16/CD32 Fc block (553142, BD Biosciences). After incubation for 5 minutes, 50 μL of an antibody cocktail in FACS buffer containing 1 μL of NeuN antibody (ab190565, Abcam) and Recombinant RNase Inhibitor was added. Samples were incubated with staining antibodies on ice for 30 minutes with gentle shaking, followed by centrifugation at 300 g for 10 minutes at 4°C. Nuclei were resuspended in 500 μL of FACS buffer containing Recombinant RNase Inhibitor, and 1 μL of Hoechst 33342 was added. NeuN-positive and NeuN-negative nuclei were sorted separately into 1.5 mL DNA LoBind tubes containing 1 mL of final buffer (200 μL UltraPure BSA, AM2618, Thermo, 800 μL PBS, and 5 μL Protector RNase Inhibitor). A total of 50,000 single nuclei were sorted per sample. Nuclei concentrations were adjusted to ∼1 million nuclei/mL.

### Chromium 10× Library Generation and Illumina Sequencing

Chromium 10× library generation and Illumina sequencing were conducted following previously published methods (Tsai AP et al, 2023). Reagents for the Chromium Single Cell 3′ Library & Gel Bead Kit v3.1 (10× Genomics, Pleasanton, USA) were thawed and prepared following the manufacturer’s protocol. The nuclei/master mix solution was adjusted to target 10,000 nuclei per sample and loaded onto a standard Chromium Controller (10× Genomics) as instructed. Library construction was carried out using the Chromium Single Cell 3′ Library Construction Kit v3.1. All reactions were performed according to the manufacturer’s protocol using the recommended reagents, consumables, and instruments. Quality control for cDNA and libraries was conducted using a Bioanalyzer (Agilent, Santa Clara, USA) at the Stanford Protein and Nucleic Acid Facility. Illumina sequencing of the 10× snRNA-seq libraries was performed by Novogene (Sacramento, USA). Multiplexed libraries were sequenced using 2×150-bp paired-end reads in a single S4 lane on an Illumina NovaSeq S4 (Illumina, San Diego, USA), targeting 100 million reads per library. Data then undergoes base-calling, demultiplexing, and FASTQ file generation to prepare for further analysis. Raw data is at: GEO repository GSE324505: GSE324504.

### snRNA-Seq Data Preprocessing and Quality Control

The CellRanger pipeline was used to align reads from each sample to the mouse mm10 (GENCODE vM23/Ensemble98) reference genome and generate expression matrices for downstream analyses. For each sample, nuclei were filtered based on their number of genes (greater than 200 but less than 5000) and their percentage of mitochondrial genes (less than 5%). Genes expressed in fewer than 3 nuclei were also removed. For each sample, counts were normalized and log-scaled, the top 2000 highly variable features were identified, and principal component analysis (PCA) was performed using the standard Seurat pipeline.

### Sample Integration and Tumor Cell Identification

We ran each sample through the copy number prediction software tools SCEVAN(De Falco A et al, 2023) and Copykat(Gao R et al, 2021) to predict which clusters represented tumor cells. Nuclei with strong expression of CD45 (immune cells) and NeuN (neurons) were used as reference non-neoplastic cells for these analyses. Based on these quality control measures, one of the MR3-treated samples was deemed poor quality (high number of nuclei removed based on the number of genes and/or identified as unreliable by SCEVAN and Copykat, so this sample was withdrawn from further analyses, leaving three untreated samples and two treated samples). Samples were then merged and integrated using the Seurat canonical correlation analysis (CCA) pipeline, and cells were clustered using 30 dimensions and a resolution of 0.5. This yielded an integrated dataset with 11,940 NeuN□ nuclei. Cluster identities were determined based on marker gene expression and copy number variation analysis, and clusters with shared cell identities were merged.

### snRNA-Seq Data Differential Expression Analysis and Pathway Enrichment Analysis

Differential expression analysis between MR3-treated and untreated cells was performed using the Wilcoxon rank-sum test implemented in the FindMarkers function in Seurat. Genes with absolute log2 fold change>0.58 and expressed in more than 10% of cells were considered significantly differentially expressed. Pathway enrichment analysis was performed separately for up- and down-regulated genes identified from single-nucleus analyses. GO enrichment was conducted using the clusterProfiler enrichGO function with the org.Mm.eg.db annotation database, considering biological process (BP), molecular function (MF), and cellular component (CC) ontologies. Kyoto Encyclopedia of Genes and Genomes (KEGG) pathway enrichment analysis was performed with the enrichKEGG function. Gene symbols were converted to Entrez IDs and mapped back to gene symbols for visualization.

### Mouse snRNA-Seq Pseudo-Bulk Construction and Differential Expression Analysis

To perform sample-aware differential expression analysis and avoid pseudoreplication, raw UMI counts were aggregated at the individual animal level to generate pseudo-bulk profiles. Specifically, counts from all nuclei belonging to the same mouse (based on orig.ident) were summed to obtain one gene-by-sample count vector per animal. This resulted in five biological replicates in total, including three untreated control mice (CN1–CN3) and two MR3-treated mice (TN2–TN3). Low-expression genes were filtered using filterByExpr in edgeR, and library sizes were normalized using the TMM method. Counts were transformed with voom and analyzed with a limma linear model with empirical Bayes moderation. Genes with an FDR-adjusted p-value < 0.05 and |log_2_ fold change| > 1 were considered significantly differentially expressed.

### Patient Information

Tissue samples were obtained from three glioma patients (P1-P3). All details are in Table S1.

### Human Sample Collection and Storage

Human tissue samples were collected from patients at Stanford Hospital after obtaining informed consent pre-operatively in accordance with an approved Stanford institutional review board-approved protocol (IRB- 12625). Three patient tissue samples from the tumor core were obtained intra-operatively and placed immediately in Dulbecco’s Modified Eagle Medium (DMEM) (Gibco^TM^) on ice before transferring to a six-well plate with either 10 μM MR3 or vehicle (DMSO) in DMEM. Samples were incubated at 37°C with 5% CO_2_ for 24 hours, then dried using Kim wipes to remove residual blood or media, and subsequently sectioned in sterile Petri dishes. All remaining samples were snap-frozen in liquid nitrogen and stored at – 80°C prior to RNA extraction. P1 had additional plain tissue that was snap frozen and profiled without treatment.

### Tissue Homogenization and RNA Isolation

Frozen tissues were pulverized in liquid nitrogen using DEPC-treated, autoclaved mortars and pestles. Approximately 0.5 mL of STAT60 (AMSBIO) was added during tissue homogenization, and the lysate was transferred to pre-chilled Eppendorf tubes. An additional 0.5 mL STAT60 was then added to obtain a final volume of 1 mL. Total RNA was isolated according to the manufacturer’s instructions using chloroform-based phase separation, and RNA was eluted in 20-40 μL of nuclease-free water, adjusted according to pellet size.

### RNA Quality Control and Storage

RNA concentration and purity were measured using a NanoDrop spectrophotometer. Samples with concentrations above 50 ng/μL and 260/280 ratios greater than 1.7 were considered to have passed quality control and were sent for sequencing. Aliquots were labeled on both cap and sidewall, sealed with parafilm, and stored at –80°C. For sequencing, samples were shipped on dry ice to Novogene (https://www.novogene.com/us-en/) with corresponding electronic and physical documentation as per vendor requirements. Raw data is at: GEO repository GSE324505: GSE322979.

### Human Bulk RNA-Seq Processing and Differential Expression Analysis

Raw RNA-seq reads were processed on the Galaxy platform (usegalaxy.org). Sequencing reads were aligned to the human reference genome hg19 using RNA STAR (Galaxy Version 2.7.11b+galaxy0), and counts per gene were obtained by running featureCounts (Galaxy Version 2.1.1+galaxy0). Gene identifiers were annotated to ENSEMBL IDs and gene symbols, respectively. Raw counts were imported into R for differential expression analysis using DESeq2. A paired design was implemented by including patient identity as a blocking factor in the DESeq2 model (∼ patient + treatment). Prior to analysis, low-expressed genes with fewer than 5 counts in at least two samples were filtered out. Filtered counts were then normalized, and variance stabilization was performed using the rlog transformation. Differential expression testing between conditions was conducted using the Wald test implemented in DESeq2. Genes with a p-value < 0.05 and log2 fold change > 1 were considered significantly differentially expressed.

### Pathway Enrichment Analysis

To gain biological insight into the identified DEGs, GO enrichment analyses were performed using the clusterProfiler package and the org.Hs.eg.db database with default settings. Enrichment was tested separately for upregulated and downregulated genes against all expressed genes as background.

### PCA and Dimensional Reduction

We applied PCA to visualize MR3-treated and untreated samples in two dimensions using the mouse pseudo-bulk RNA-seq data.

### Cross-Species Transcriptomic Datasets Integration

Three datasets were integrated to identify conserved MR3-rescued glioma dysregulated genes: 1. TCGA/GTEx GBM versus normal brain, 2. MR3-treated versus untreated GL261 mouse tumors (pseudo-bulk RNA-seq and snRNA-seq), 3. MR3-treated versus untreated human glioma samples (bulk RNA-seq). For the mouse dataset, we generated pseudo-bulk profiles from snRNA-seq data and compared MR3-treated tumors with untreated tumor samples. For the human dataset, we compared bulk RNA-seq profiles between MR3-treated and untreated tumor samples. In both the mouse and human MR3 datasets, we only kept genes with one-to-one orthologs in the other species, ensuring that the results were directly comparable across species. When integrating the three datasets to identify overlapping gene sets, we limited the analysis to protein-coding genes that were present in all datasets.

### Cell-Cell Communication Analysis

To investigate intercellular communication among immune and tumor cell populations, we applied the CellChat R package (v2.2.0) to our snRNA-seq data. Processed snRNA-seq data were first imported from Seurat objects, with subcluster annotations as cell identities. Separate CellChat objects were created for MR3□ (untreated) and MR3□ (treated) groups using the function createCellChat. Intercellular communications were identified based on the CellChatDB.mouse database. Overexpressed signaling genes were detected using identifyOverExpressedGenes, and significant ligand–receptor interactions were identified by identifyOverExpressedInteractions with default settings. Communication probabilities between subclusters were computed using the computeCommunProb function in CellChat with population.size = TRUE. A truncatedMean approach (trim = 0.13) was applied to reduce the effect of extreme expression values. Weak or nonsignificant interactions were removed using filterCommunication and requiring at least 20 cells per group (min.cells = 20). Pathway-level communication probabilities were inferred using computeCommunProbPathway, and the overall network was summarized with aggregateNet with default settings.

### Web Platform and Data Publication

The MiroScape website was built using front-end architecture with the React framework. React-plotly.js was used for data visualization. The platform is openly available and deployed through GitHub Pages. snRNA-seq data are displayed in an interactive interface built with ShinyCell2 and deployed through shiny.io. The interface is embedded into the website to provide direct and uninterrupted access for users.

### Computational Reproducibility

Computational analyses were performed in R (v4.4.2) and Python (v3.13.3). snRNA-seq data were processed with Seurat (v5.3.0). Pseudobulk differential expression was conducted with the edgeR (v4.4.2)– limma (v3.62.2) workflow, and human bulk RNA-seq differential expression was performed with DESeq2 (v1.40.2) using a paired design with patient as a blocking factor. Cell–cell communication analysis was conducted with CellChat (v2.2.0), and pathway enrichment with clusterProfiler (v4.8.1). Bulk RNA-seq reads were aligned with STAR and quantified using featureCounts.

## Data Availability

Processed data and analysis are available for download on our interactive platform, MiroScape, under the MiroScripts/Source Data tab. All original code is available on GitHub at https://github.com/miroscape/MiroScape. Raw sequencing data are at GEO repository GSE324505: GSE324504, GSE322979. TCGA (Weinstein JN et al, 2013) glioma samples and non-tumor brain cortex from the GTEx (The genotype-tissue expression (gtex) project, 2013) dataset were analyzed with GEPIA3 (Kang YJ et al, 2025) (https://gepia3.bioinfoliu.com/). Any additional information required to reanalyze the data reported in this paper is available from Xinnan Wang.

## Acknowledgement

We thank E.R. Lopez, M. Atkins, E.K. Costa, Drs., J. Qiu, A.G. Sainz, J. Haberberger, A. Isakova, C. Jackson, and C.K. Petritsch for technical support and discussion; the following funders: National Institutes of Health (RO1NS128040 and RO1GM143258, X.W.), Stanford Wu Tsai Neurosciences Institute Translate Award (X.W. and M.L.).

## Author Contributions

X.W. conceived and supervised the project. M.L. supervised overall patient tumor collection and mouse model setup. Z.D. and M.-H.L. analyzed transcriptomic data and performed visualization. Z.D. built the MiroScape platform. B.H.B. and A.P.T. preprocessed mouse snRNA-seq data. K.C. established the mouse model. A.P.T., K.C., and T.W.-C. assisted with snRNA-seq sample preparation. L.H.K., J.C., and G.L. coordinated patient recruitment, consent, and surgery. J.C. processed patient samples. Z.D. and X.W. wrote the manuscript with input from all authors. Some figures were created with https://BioRender.com/w9trdi8.

## Supplementary Figure Legends

**Figure S1. Quality Control of Our Datasets.**

**(A)** Quality control metrics for mouse snRNA-seq by sample. For each sample, metrics show as: nCount_RNA (left), nFeature_RNA (middle), and mitochondrial transcript percentage (right). From top left to bottom right: CN1, CN2, CN3, TN1, TN2 and TN3.

**(B)** Quality control metrics for mouse snRNA-seq by cell type: nCount_RNA (left), nFeature_RNA (middle), and mitochondrial transcript percentage (right).

**(C)** Tumor annotation and quality control. Top: Proportions of filtered, normal, and tumor cells across annotated cell types by SCEVAN. Middle: CopyKAT-inferred ploidy status. Bottom: Cell cycle phase distribution.

**Figure S2. Overall Analysis of Our Mouse Data.**

**(A)** Expression of marker genes for major cell types.

**(B)** Top differentially expressed genes across major cell types.

**(C)** PCA of pseudo-bulked mouse snRNA-seq data using all detected genes.

**(D)** Cross-species overlap of MR3-regulated genes. Left: Venn diagram of upregulated genes in MR3-treated mouse and human tumors. Right: Venn diagram of downregulated genes.

**(E)** Cell type proportions per mouse in untreated control (CN) and MR3-treated (TN) groups.

**(F)** Tumor proportion per mouse in untreated control (CN) and MR3-treated (TN) groups.

**Figure S3. Further Analysis of Our Mouse Data.**

**(A)** Volcano plots comparing differential gene expression in macrophages treated with and without MR3. DE: differential expression, identified using Seurat’s FindAllMarkers function based on single-cell–level analysis.

**(B-C)** Functional enrichment analysis of DEGs in macrophages before (MR3□) and after (MR3□) treatment. Both GO (Biological Process, BP; Cellular Component, CC; and Molecular Function, MF categories) and KEGG pathway analyses were performed. (B) Top 5 significantly upregulated pathways; (C) Top 5 significantly downregulated pathways.

Volcano plots comparing differential gene expression in macrophages treated with and without MR3. Differential expression analysis was performed using a sample-aware, pseudobulk approach, in which gene counts were aggregated at the biological replicate level prior to statistical testing.

**(E-F)** Functional enrichment analysis of DEGs in macrophages before (MR3□) and after (MR3□) treatment. GO (Biological Process, BP; Cellular Component, CC; and Molecular Function, MF categories) analysis was performed. (E) Top significantly upregulated pathways; (F) Top significantly downregulated pathways.

**(G)** UMAP projection of macrophage, microglia, lymphoid, and tumor cells in MR3-treated and untreated groups.

**Figure S4. Subclustering Analysis of Our Mouse Data.**

**(A)** UMAP projection showing the expression of canonical marker genes for microglia, macrophages, and DCs. The color intensity represents the normalized expression level of each marker gene across cells.

**(B)** Dot plot showing the expression of the top 3 DEGs in macrophages, microglia, and DCs. The color intensity of each dot represents the average expression level, while the dot size indicates the proportion of cells expressing the corresponding gene within each cluster.

**(C)** UMAP projection showing the expression of canonical marker genes for NK cells, regulatory-like CD4□ T cells, and CD8□ T cells. The color intensity represents the normalized expression level of each marker gene across cells.

**(D)** Dot plot showing the expression of the top 3 DEGs in NK cells, regulatory-like CD4□ T cells, and CD8□ T cells. The color intensity of each dot represents the average expression level, while the dot size indicates the proportion of cells expressing the corresponding gene within each cluster.

**(E)** UMAP projections showing tumor subtype annotations.

**(F)** Dot plot showing the expression of the top 3 DEGs in tumor subclusters. The color intensity of each dot represents the average expression level, while the dot size indicates the proportion of cells expressing the corresponding gene within each cluster.

**(G)** Bar plots showing the relative proportions of immune cell subclusters before (control) and after MR3 treatment.

**(H)** Bar plots showing the relative proportions of tumor cell subclusters before (control) and after MR3 treatment.

**Figure S5. Cell-Cell Communication Analysis.**

**(A)** Bar plots comparing the number of inferred interactions (left) and interaction strength (right) between MR3□ and MR3□ conditions.

**(B-E)** Heatmap visualization of CellChat-inferred cell-cell communication in MR3□ (B, D) and MR3□ (C, E) conditions, with color intensity indicating either the number of interactions (B, C) or the interaction strength (D, E).

**(F, G)** Dot plots depicting CellChat analysis of cell-cell communication in the MR3□ (F) and MR3□ (G) conditions, with MAC1 as the signal receiver. Dot size represents significance, and color indicates communication probability.

**Figure S6. T Cell Activation and Mini-Screen of MIRO1-Relevant Pathways.**

**(A)** Dot plots illustrating the expression of representative CD8^+^ T cell activation-, cytotoxicity- and interferon response–related genes in CD8^+^ T cells from control (CD8^+^ T-) and MR3-treated (CD8^+^ T+) groups. Dot size indicates the fraction of CD8^+^ T cells expressing each gene, and color intensity corresponds to the scaled average expression level.

**(B-D)** Dot plots illustrating the expression of selected pathway-related genes across myeloid subclusters in MR3□ and MR3□ conditions. The size of each dot represents the proportion of cells expressing the gene, and the color intensity indicates the average expression level. Subclusters with fewer than 20 cells were excluded.

**(B)** NF-κB pathway–related genes.

**(C)** Genes involved in mitochondrial motility, dynamics, and mitophagy.

**(D)** Genes related to the cGAS–STING pathway and mtDNA release.

## Supplementary Table Title

**Table S1. Patient and Sample Information.**

