## Supplementary Figure 1-6 and Table for "Cross-Species Transcriptomic Integration Reveals a MIRO1-Mediated Macrophage–T Cell Axis in Glioma"

**Fig. S6. T Cell Activation and Mini-Screen of MIRO1-Relevant Pathways.**
**(A)** Dot plots illustrating the expression of representative $CD8^{+}$ T cell activation-, cytotoxicity- and interferon response–related genes in $CD8^{+}$ T cells from control ($CD8^{+}$ T-) and MR3-treated ($CD8^{+}$ T+) groups. Dot size indicates the fraction of $CD8^{+}$ T cells expressing each gene, and color intensity corresponds to the scaled average expression level.
**(B-D)** Dot plots illustrating the expression of selected pathway-related genes across myeloid subclusters in $MR3^{-}$ and $MR3^{+}$ conditions. The size of each dot represents the proportion of cells expressing the gene, and the color intensity indicates the average expression level. Subclusters with fewer than 20 cells were excluded.
**(B)** NF-κB pathway–related genes.
**(C)** Genes involved in mitochondrial motility, dynamics, and mitophagy.
**(D)** Genes related to the cGAS–STING pathway and mtDNA release.

Fig S1

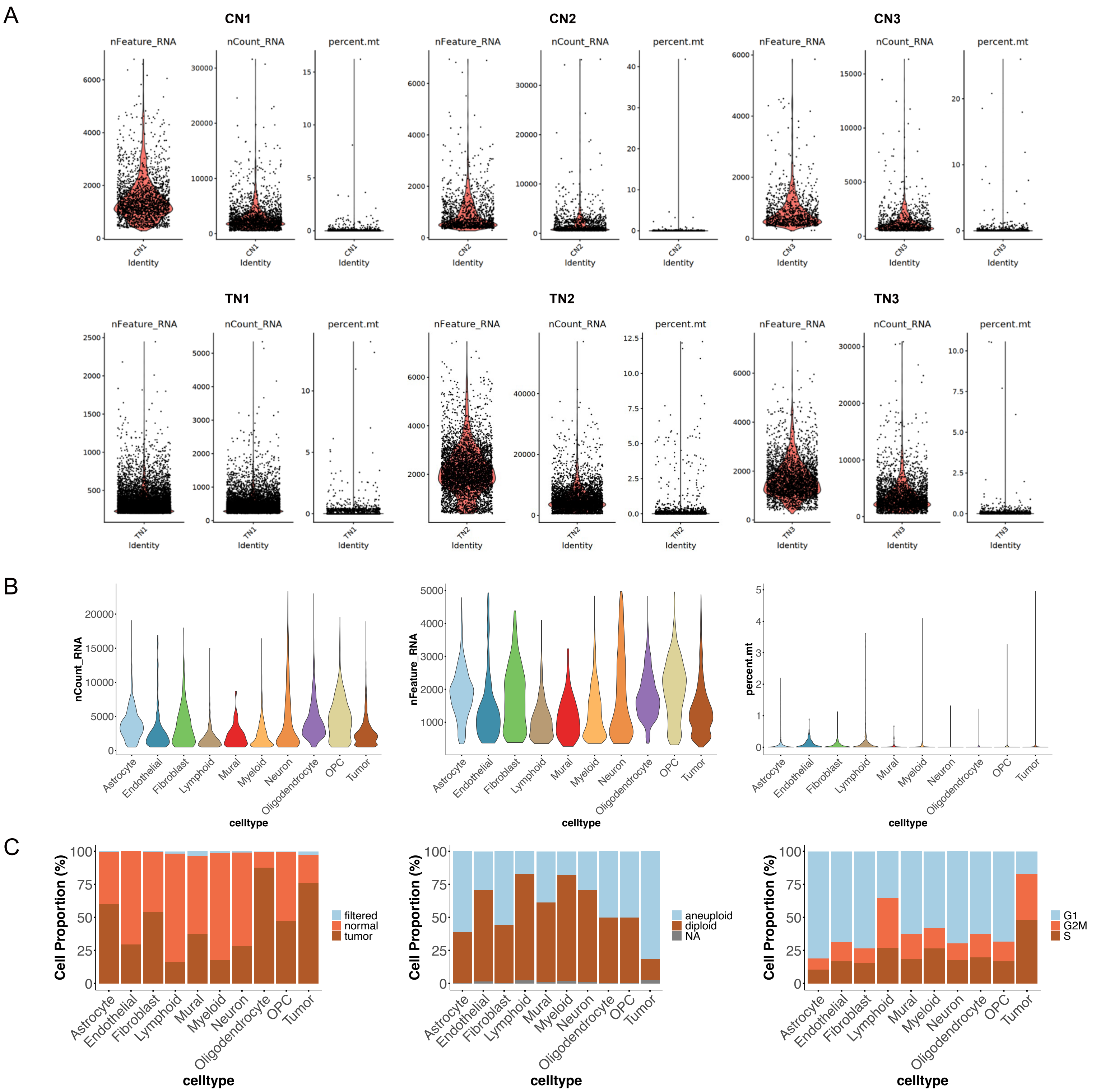
A
CN1
CN2
CN3
TN1
TN2
TN3
nFeature_RNA
nCount_RNA
percent.mt
Identity
B
nCount_RNA
nFeature_RNA
percent.mt
celltype
Astrocyte
Endothelial
Fibroblast
Lymphoid
Mural
Myeloid
Neuron
Oligodendrocyte
OPC
Tumor
C
Cell Proportion (%)
filtered
normal
tumor
aneuploid
diploid
NA
G1
G2M
S
celltype

Fig S2

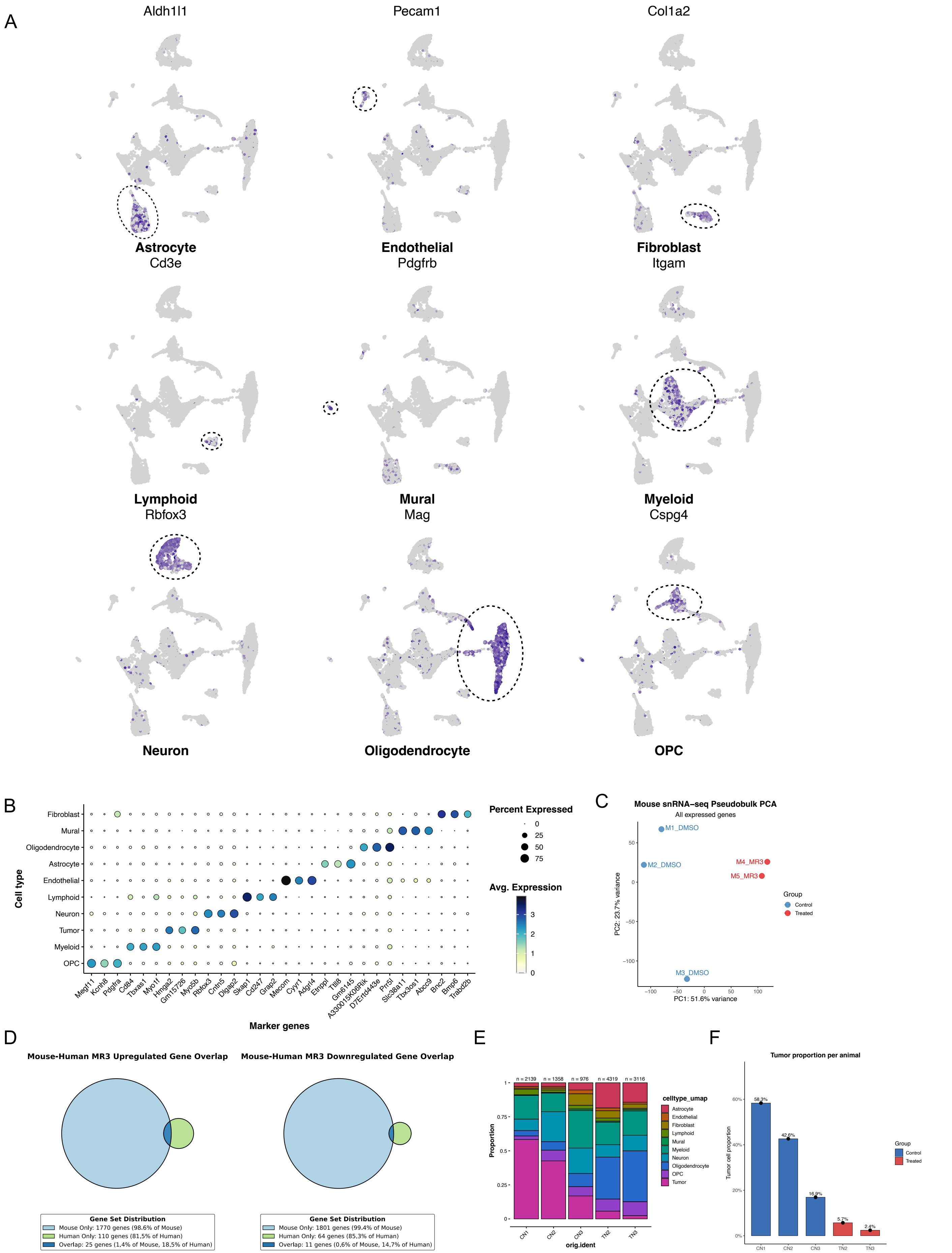
A
Aldh1l1
Pecam1
Col1a2
Astrocyte
Cd3e
Endothelial
Pdgfrb
Fibroblast
Itgam
Lymphoid
Rbfox3
Mural
Mag
Myeloid
Cspg4
Neuron
Oligodendrocyte
OPC
B
Fibroblast
Mural
Oligodendrocyte
Astrocyte
Endothelial
Lymphoid
Neuron
Tumor
Myeloid
OPC
Cell type
Megf11 Kcnh8 Pdgfra Cd84 Tbxas1 Myo1f Hmga2 Gm15726 Myo5b Rbfox3 Cntn5 Dlgap2 Skap1 Cd247 Grap2 Mecom Cyyr1 Adgrl4 Etnppl Tlll6 Gm6145 A330015K06Rik D7Ertd443e Prr5l Slc38a11 Tbx3os1 Abcc9 Bnc2 Bmp6 Trabd2b
Marker genes
Percent Expressed
0
25
50
75
Avg. Expression
3
2
1
0
C
Mouse snRNA−seq Pseudobulk PCA
All expressed genes
M1_DMSO
M2_DMSO
M4_MR3
M5_MR3
M3_DMSO
Group
Control
Treated
PC2: 23.7% variance
PC1: 51.6% variance
D
Mouse-Human MR3 Upregulated Gene Overlap
Mouse-Human MR3 Downregulated Gene Overlap
Gene Set Distribution
Mouse Only: 1770 genes (98.6% of Mouse)
Human Only: 110 genes (81.5% of Human)
Overlap: 25 genes (1.4% of Mouse, 18.5% of Human)
Gene Set Distribution
Mouse Only: 1801 genes (99.4% of Mouse)
Human Only: 64 genes (85.3% of Human)
Overlap: 11 genes (0.6% of Mouse, 14.7% of Human)
E
n = 2139
n = 1358
n = 976
n = 4319
n = 3116
Proportion
CN1
CN2
CN3
TN2
TN3
orig.ident
celltype_umap
Astrocyte
Endothelial
Fibroblast
Lymphoid
Mural
Myeloid
Neuron
Oligodendrocyte
OPC
Tumor
F
Tumor proportion per animal
Tumor cell proportion
58.3%
42.6%
16.9%
5.7%
2.4%
CN1
CN2
CN3
TN2
TN3
Group
Control
Treated

Fig S3

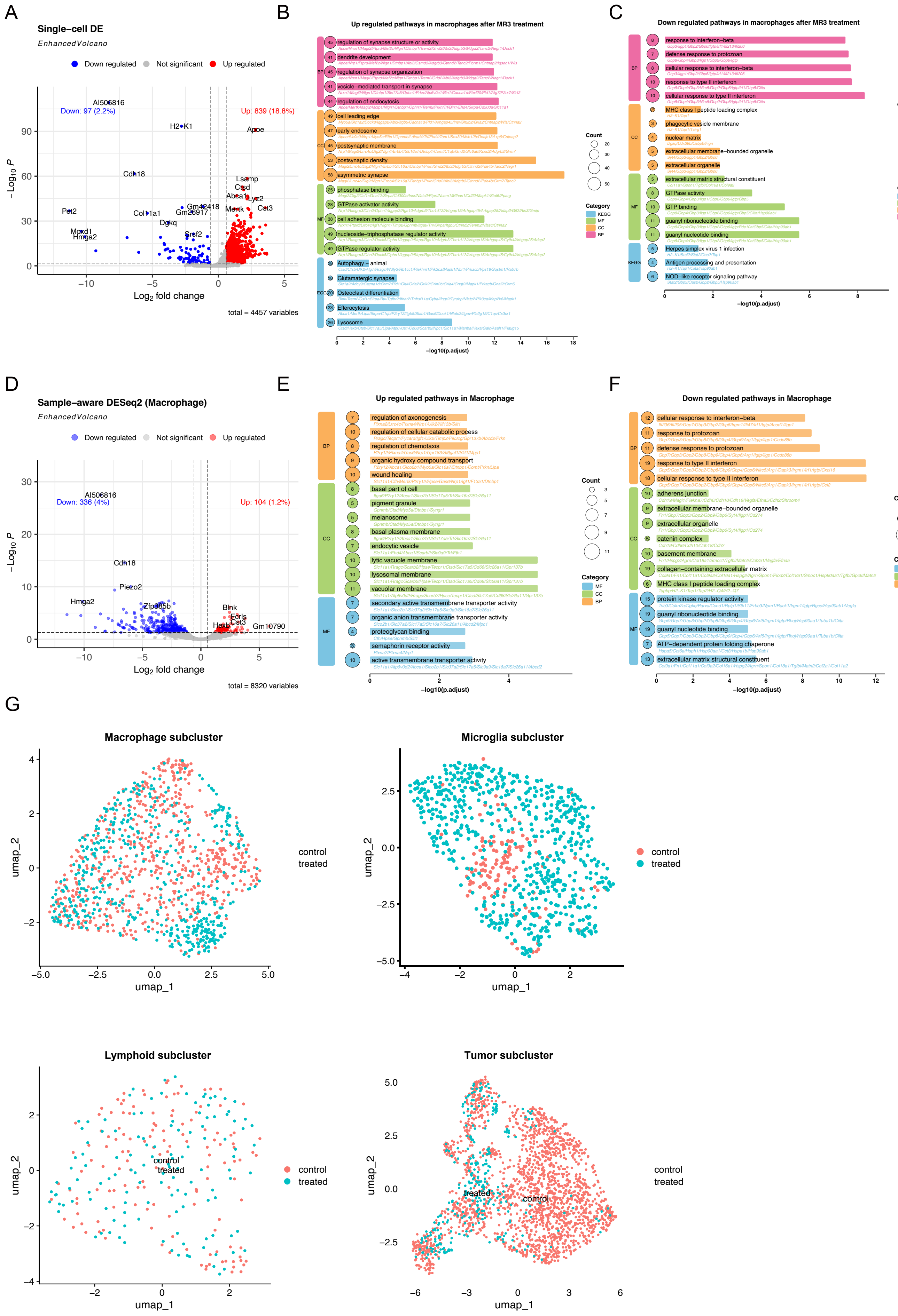
A
Single−cell DE
EnhancedVolcano
Down regulated
Not significant
Up regulated
Down: 97 (2.2%)
Up: 839 (18.8%)
AI506816
H2−K1
Apoe
Cdh18
Lsamp
Ctsd
Abca1
Lyz2
Mertk
Cst3
Pet2
Col11a1
Gm26917
Gm42418
Dgkq
Mcrd1
Hmga2
Srgf2
− Log10 P
Log2 fold change
total = 4457 variables
B
Up regulated pathways in macrophages after MR3 treatment
regulation of synapse structure or activity
dendrite development
regulation of synapse organization
vesicle−mediated transport in synapse
regulation of endocytosis
cell leading edge
early endosome
postsynaptic membrane
postsynaptic density
asymmetric synapse
phosphatase binding
GTPase activator activity
cell adhesion molecule binding
nucleoside−triphosphatase regulator activity
GTPase regulator activity
Autophagy − animal
Glutamatergic synapse
Osteoclast differentiation
Efferocytosis
Lysosome
Count
Category
KEGG
MF
CC
BP
−log10(p.adjust)
C
Down regulated pathways in macrophages after MR3 treatment
response to interferon−beta
defense response to protozoan
cellular response to interferon−beta
response to type II interferon
cellular response to type II interferon
MHC class I peptide loading complex
phagocytic vesicle membrane
nuclear matrix
extracellular membrane−bounded organelle
extracellular organelle
extracellular matrix structural constituent
GTPase activity
GTP binding
guanyl ribonucleotide binding
guanyl nucleotide binding
Herpes simplex virus 1 infection
Antigen processing and presentation
NOD−like receptor signaling pathway
Count
Category
KEGG
MF
CC
BP
−log10(p.adjust)
D
Sample−aware DESeq2 (Macrophage)
EnhancedVolcano
Down regulated
Not significant
Up regulated
Down: 336 (4%)
Up: 104 (1.2%)
AI506816
Cdh18
Piezo2
Hmga2
Zfp385b
Blnk
Fcrls
Cst3
Hexb
Gm10790
− Log10 P
Log2 fold change
total = 8320 variables
E
Up regulated pathways in Macrophage
regulation of axonogenesis
regulation of cellular catabolic process
regulation of chemotaxis
organic hydroxy compound transport
wound healing
basal part of cell
pigment granule
melanosome
basal plasma membrane
endocytic vesicle
lytic vacuole membrane
lysosomal membrane
vacuolar membrane
secondary active transmembrane transporter activity
organic anion transmembrane transporter activity
proteoglycan binding
semaphorin receptor activity
active transmembrane transporter activity
Count
Category
MF
CC
BP
−log10(p.adjust)
F
Down regulated pathways in Macrophage
cellular response to interferon−beta
response to protozoan
defense response to protozoan
response to type II interferon
cellular response to type II interferon
adherens junction
extracellular membrane−bounded organelle
extracellular organelle
catenin complex
basement membrane
collagen−containing extracellular matrix
MHC class I peptide loading complex
protein kinase regulator activity
guanyl ribonucleotide binding
guanyl nucleotide binding
ATP−dependent protein folding chaperone
extracellular matrix structural constituent
Count
Category
MF
CC
BP
−log10(p.adjust)
G
Macrophage subcluster
umap_2
umap_1
control
treated
Microglia subcluster
umap_2
umap_1
control
treated
Lymphoid subcluster
umap_2
umap_1
control
treated
Tumor subcluster
umap_2
umap_1
treated
control
control
treated

### Fig S4

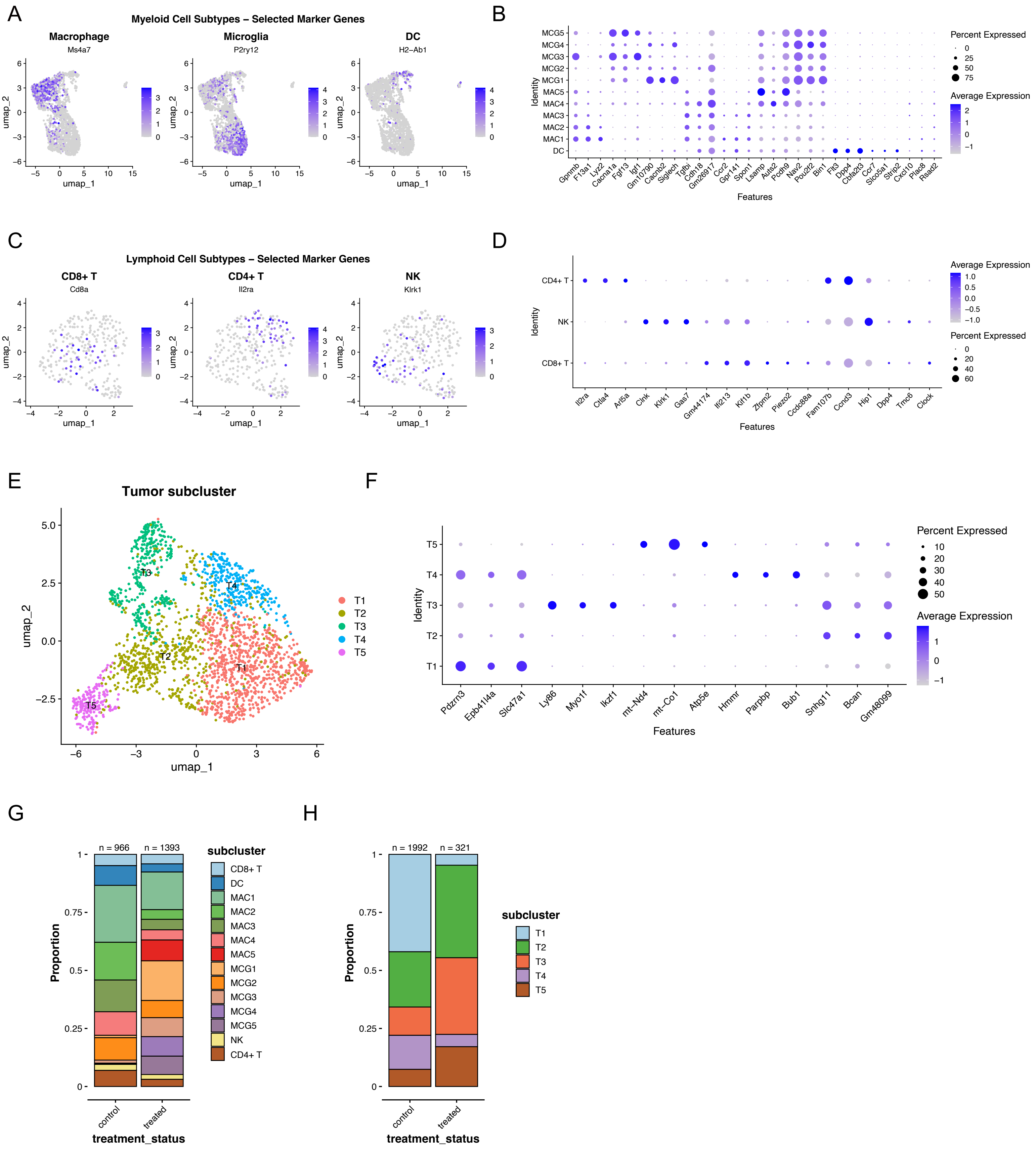
A
Myeloid Cell Subtypes – Selected Marker Genes
Macrophage
Ms4a7
Microglia
P2ry12
DC
H2-Ab1
umap_1
umap_2
B
Percent Expressed
Average Expression
Identity
Features
C
Lymphoid Cell Subtypes – Selected Marker Genes
CD8+ T
Cd8a
CD4+ T
Il2ra
NK
Klrk1
D
E
Tumor subcluster
F
G
H
subcluster
Proportion
treatment_status
control
treated
n = 966
n = 1393
n = 1992
n = 321

### Fig S5

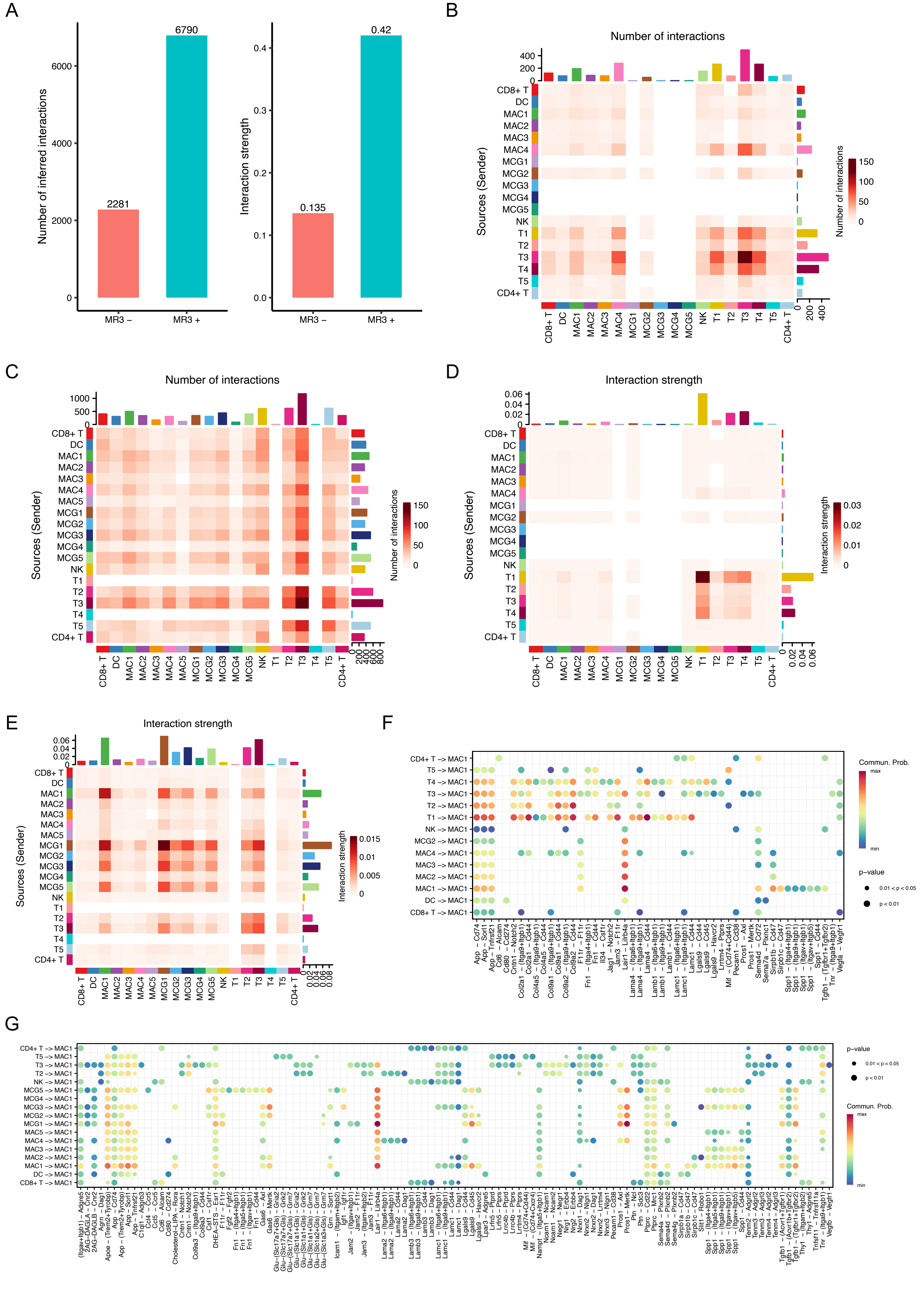

A
Number of inferred interactions
2281
6790
MR3 –
MR3 +
Interaction strength
0.135
0.42
B
Number of interactions
Sources (Sender)
CD8+ T
DC
MAC1
MAC2
MAC3
MAC4
MCG1
MCG2
MCG3
MCG4
MCG5
NK
T1
T2
T3
T4
T5
CD4+ T
Number of interactions
C
Number of interactions
Sources (Sender)
MAC5
D
Interaction strength
Sources (Sender)
Interaction strength
E
Interaction strength
Sources (Sender)
Interaction strength
F
CD4+ T -> MAC1
T5 -> MAC1
T4 -> MAC1
T3 -> MAC1
T2 -> MAC1
T1 -> MAC1
NK -> MAC1
MCG2 -> MAC1
MAC4 -> MAC1
MAC3 -> MAC1
MAC2 -> MAC1
MAC1 -> MAC1
DC -> MAC1
CD8+ T -> MAC1
Commun. Prob.
max
min
p-value
0.01 < p < 0.05
p < 0.01
G
MCG5 -> MAC1
MCG4 -> MAC1
MCG3 -> MAC1
MCG2 -> MAC1
MCG1 -> MAC1
MAC5 -> MAC1
p-value
Commun. Prob.

Fig S6

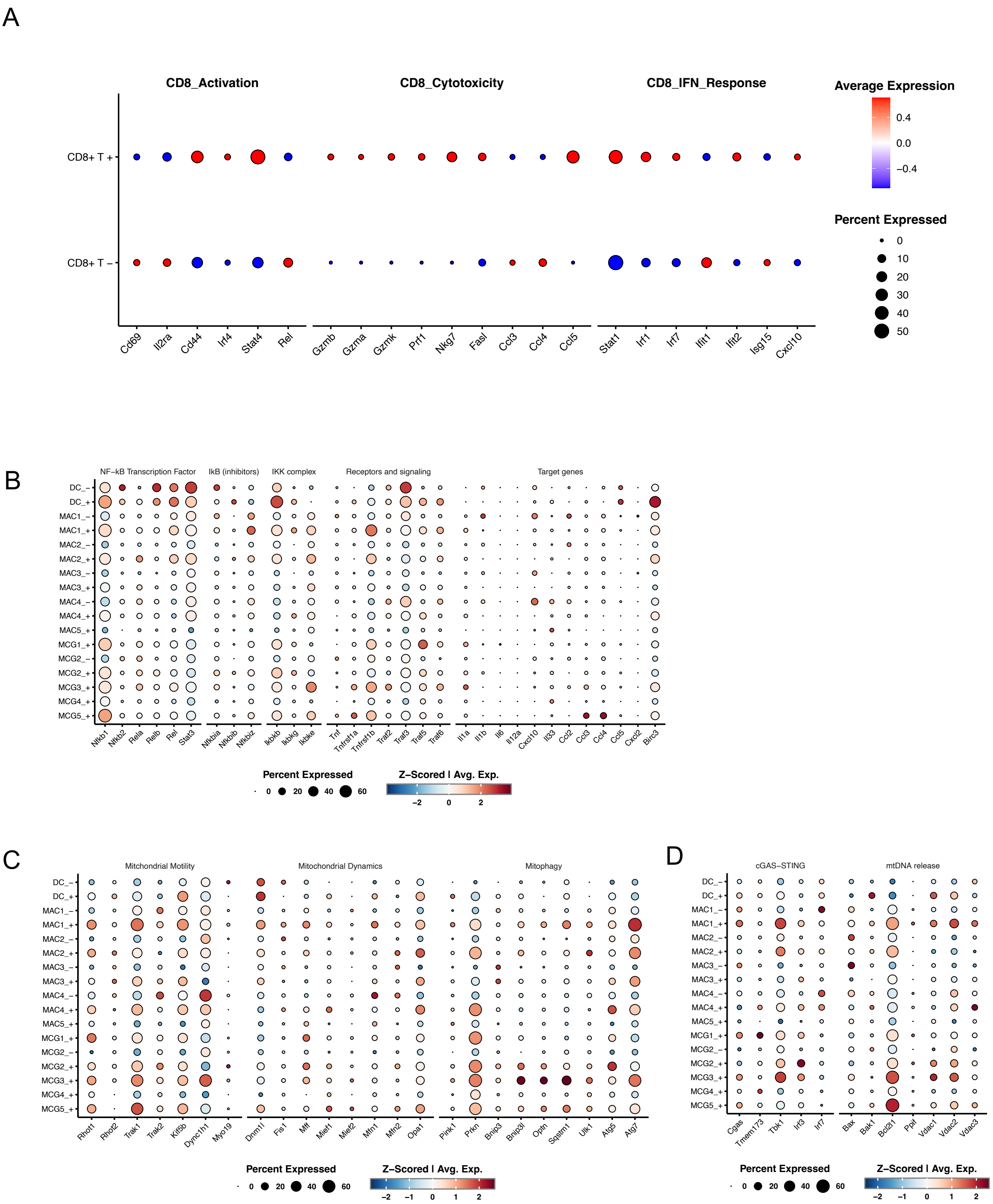
A
CD8_Activation
CD8_Cytotoxicity
CD8_IFN_Response
Average Expression
0.4
0.0
−0.4
Percent Expressed
0
10
20
30
40
50
CD8+ T +
CD8+ T −
Cd69
Il2ra
Cd44
Irf4
Stat4
Rel
Gzmb
Gzma
Gzmk
Prf1
Nkg7
Fasl
Ccl3
Ccl4
Ccl5
Stat1
Irf1
Irf7
Ifit1
Ifit2
Isg15
Cxcl10
B
NF−kB Transcription Factor
IkB (inhibitors)
IKK complex
Receptors and signaling
Target genes
DC_−
DC_+
MAC1_−
MAC1_+
MAC2_−
MAC2_+
MAC3_−
MAC3_+
MAC4_−
MAC4_+
MAC5_+
MCG1_+
MCG2_−
MCG2_+
MCG3_+
MCG4_+
MCG5_+
Nfkb1
Nfkb2
Rela
Relb
Rel
Stat3
Nfkbia
Nfkbib
Nfkbiz
Ikbkb
Ikbkg
Ikbke
Tnf
Tnfrsf1a
Tnfrsf1b
Traf2
Traf3
Traf5
Traf6
Il1a
Il1b
Il6
Il12a
Cxcl10
Il33
Ccl2
Ccl3
Ccl4
Ccl5
Cxcl2
Birc3
Percent Expressed
0
20
40
60
Z−Scored | Avg. Exp.
−2
0
2
C
Mitchondrial Motility
Mitochondrial Dynamics
Mitophagy
Rhot1
Rhot2
Trak1
Trak2
Kif5b
Dync1h1
Myo19
Dnm1l
Fis1
Mff
Mief1
Mief2
Mfn1
Mfn2
Opa1
Pink1
Prkn
Bnip3
Bnip3l
Optn
Sqstm1
Ulk1
Atg5
Atg7
Percent Expressed
0
20
40
60
Z−Scored | Avg. Exp.
−2
−1
0
1
2
D
cGAS−STING
mtDNA release
Cgas
Tmem173
Tbk1
Irf3
Irf7
Bax
Bak1
Bcl2l1
Ppif
Vdac1
Vdac2
Vdac3
Percent Expressed
0
20
40
60
Z−Scored | Avg. Exp.
−2
−1
0
1
2

| Patient | Date | Sex | Age | Final Path | Sample |
|---|---|---|---|---|---|
| P1 | 4/30/2024 | M | 50 | IDH-mutant grade 2 oligodendroglioma;1p/19q-codeleted | P1_Plain, P1_DMSO, P1_MR3 |
| P2 | 8/26/2024 | F | 29 | IDH-mutant grade 2 astrocytoma; P53 overexpressed; ATRX lost; GFAP positive | P2_DMSO,P2_MR3 |
| P3 | 8/27/2024 | F | 51 | IDH-wildtype grade 4 glioblastoma; ATRX intact; P53 overexpressed | P3_DMSO,P3_MR3 |

**Table S1** **Patient and Sample Information.**
